# Binding of Ca^2+^-Independent C2 Domains to Lipid Membranes: a Multi-Scale Molecular Dynamics Study

**DOI:** 10.1101/2020.10.30.361964

**Authors:** Andreas Haahr Larsen, Mark S.P. Sansom

**Affiliations:** Department of Biochemistry, University of Oxford, South Parks Road, OX1 3QU, Oxford, UK

## Abstract

C2 domains facilitate protein-lipid interaction in cellular recognition and signalling processes. They possess a β-sandwich structure, with either type I or type II topology. C2 domains can interact with anionic lipid bilayers in either a Ca2+-dependent or a Ca2+-independent manner. The mechanism of recognition of anionic lipids by Ca2+-independent C2 domains is incompletely understood. We have used molecular dynamics (MD) simulations to explore the membrane interactions of six Ca2+– independent C2 domains, from KIBRA, PI3KC2α, RIM2, PTEN, SHIP2, and Smurf2. In coarse grained MD simulations these C2 domains bound to lipid bilayers, forming transient interactions with zwitterionic (phosphatidylcholine, PC) bilayers compared to long lived interactions with anionic bilayers also containing either phosphatidylserine (PS) or PS and phosphatidylinositol bisphosphate (PIP2). Type I C2 domains bound non-canonically via the front, back or side of the β sandwich, whereas type II C2 domains bound canonically, via the top loops (as is typically the case for Ca2+-dependent C2 domains). C2 domains interacted strongly (up to 120 kJ/mol) with membranes containing PIP2 causing the bound anionic lipids to clustered around the protein. The C2 domains bound less strongly to anionic membranes without PIP2 (<50 kJ/mol), and most weakly to neutral membranes (<33 kJ/mol). Productive binding modes were identified and further analysed in atomistic simulations. For PTEN and SHIP2, CG simulations were also performed of the intact enzymes (i.e. phosphatase domain plus C2 domain) with PIP2-contating bilayers and the roles of the two domains in membrane localization were compared. From a methodological perspective, these studies establish a multiscale simulation protocol for studying membrane binding/recognition proteins, capable of revealing binding modes alongside details of lipid binding affinity and specificity.

## Introduction

Lipid specific membrane recognition plays a key role in the biology of eukaryotic cells. A number of families of recognition domains exist. Of especial importance are domains that recognize phosphoinositide (PI) lipids (Kutateladze, 2010; Stahelin et al., 2014), with C2 and PH domains being the most abundant families (Katan and Allen, 1999). C2 domains can enable both protein-lipid and protein-protein interactions (Corbalan-Garcia and Gómez-Fernández, 2014; Nalefski and Falke, 1996; Zhang and Aravind, 2010), and are key players in a number of cellular signalling processes involving, e.g., ubiquitination (Wiesner et al., 2007) or (de)phosphorylation (Chen et al., 2018; Gericke et al., 2013). C2 domains bind membranes and/or other proteins and thereby brings adjacent active domains in contact with their interaction partners. One intensively studied example of a C2 domain-containing protein is the phosphatase and tensin homologue (PTEN), which consists of a C2 domain and a phosphatase domain (Lee et al., 1999; Maehama and Dixon, 1999; Zhao et al., 2004). In PTEN, C2 provides a lipid-mediated anchor to the membrane, so the phosphatase domain can dephosphorylate phosphatidylinositol triphosphate (PIP_3_) to phosphatidylinositol di-3,4-phosphate (PI(3,4)P_2_). This dephosphorylation downregulates the Akt pathway, and PTEN is thus a tumour suppressor.

All C2 domains have a similar structure (Figure 1). They are composed of around 130 residues arranged as 8 β antiparallel strands (β1-8) in two β-sheets, forming a β sandwich. C2 domains have two topologies, type I or type II, which differ by a circular permutation (Corbalan-Garcia and Gómez-Fernández, 2014; Nalefski and Falke, 1996). Calcium ions control the function of many C2-domains, via Ca^2+^-mediated binding and unbinding to lipid membranes (Corbalan-Garcia and Gómez-Fernández, 2014). However, there are a group of Ca^2+^-independent C2 domains with either no or little Ca^2+^-dependency. These are the focus of the current study.

**Figure 1.**
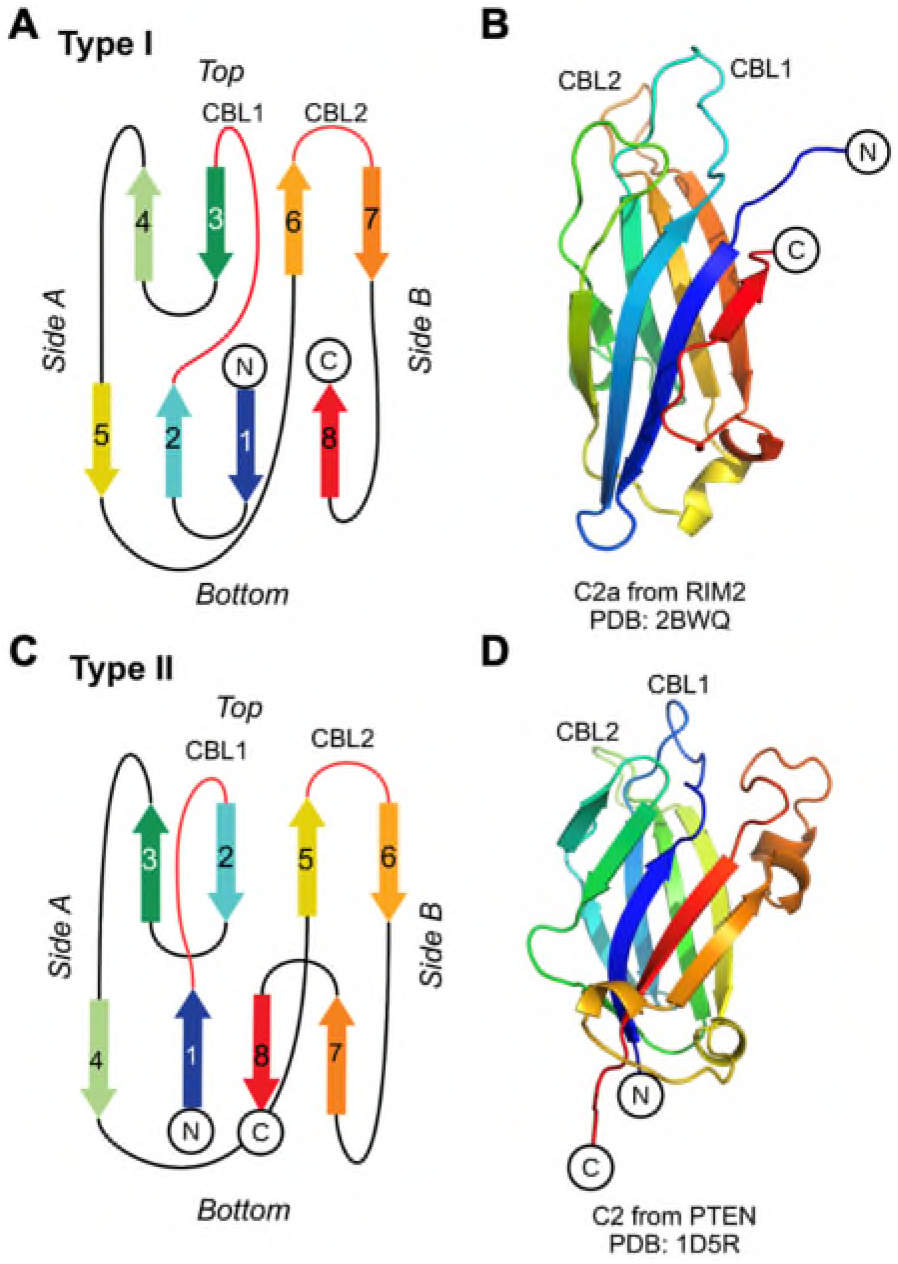
C2 structures and topologies. **A, C** Schematic diagrams of the topologies of Type I and II C domains, with β-strands numbered and connected with loops, coloured using a rainbow spectrum, from blue to red. The β-sheet containing the N- and C-termini is here denoted the “front”, and the other β-sheet the “back”. **B, D** Examples of C2 domains of each topology: the type I example is C2a from RIM2 (PDB ID 2BWQ) and the type II example is C2 from PTEN (PDB ID ID5R).

We examine how Ca^2+^-independent C2 domains interact with lipid membranes, especially those containing PI lipids. Molecular dynamics (MD) simulations have emerged as a powerful tool to examine the interactions of membrane proteins with lipids (Corradi et al., 2019; Hedger and Sansom, 2016; Manna et al., 2019) and have previously been applied to investigate the interactions of peripheral membrane proteins, especially PH domains, with PI-containing lipid bilayers (Kalli and Sansom, 2014; Lai et al., 2013, 2010; Naughton et al., 2016; Pant and Tajkhorshid, 2020; Yamamoto et al., 2016). There have been a number of MD studies of the interaction of Ca^2+^-dependent C2 domains with membranes, revealing how Ca^2+^ ions mediate interactions between the protein and anionic lipids such as phosphatidyl serine (PS) and phosphatidyl inositol phosphates (PIPs) (Alwarawrah and Wereszczynski, 2017; Banci et al., 2002; Chon et al., 2015; Jaud et al., 2007; Lai et al., 2010; Manna et al., 2008; Michaeli et al., 2017; Vermaas and Tajkhorshid, 2017). However, there have not been many simulation studies of the interactions of Ca^2+^-independent C2 domains with membranes (see e.g. (Scott et al., 2020)) other than for PTEN (S. Shenoy et al., 2012; Treece et al., 2020) despite the growing recognition of the importance of such interactions in a number of cellular processes (Stahelin et al., 2014). Here we use coarse-grained molecular dynamics (CG-MD) (Ingólfsson et al., 2014; Marrink and Tieleman, 2013) to examine and compare the interactions of six different Ca^2+^-independent C2 domains with lipid bilayers (C2 domains from KIBRA (PDB ID 6FJD), PI3KC2α (PDB ID 6BU0), RIM2 (PDB ID 2BWQ), PTEN (PDB ID 1D5R), Smurf2 (PDB ID 50KM) and SHIP2 (PDB ID 2JQZ)). We explore both their lipid specificity and the energetics of interaction. The most favourable interaction mode(s) for each C2 domain are refined by atomistic molecular dynamics (AT-MD) simulations. Based on ~1 msec total of CG simulation data for 6 C2 domains and 3 membranes, we demonstrate that the binding orientations is related to topology, and we show that the C2 domains only remain bound to negatively charged membranes and can induce clustering of multiple phosphatidylinositol-diphosphates. Our studies also provide a protocol for simulation studies of the specificity and energetics of lipid bilayer interactions of other families of membrane binding/recognition proteins.

## Results

### A protocol for determining and analysing C2 domain binding modes

To explore possible binding modes of C2 domains to lipid bilayers without bias from the initial simulation system configuration, we initiated CG simulations with the C2 domain positioned at sufficient distance that it could not “feel” the membrane, i.e. at a distance greater than the cutoff distances of the CG force field. For each simulation of an ensemble the protein molecule was rotated through a random angle before the simulation was started (Figure 2). Each simulation was run for 2 μs which proved long enough for the protein to encounter and interact with the bilayer, in some cases (dependent on the bilayer lipid composition) being able to dissociate and rebind. To ensure adequate sampling, based on previous experience with PH domains (Yamamoto et al., 2016) and other membrane proteins (e.g. phosphatidylinositol phosphate kinase PIP5K1A (Amos et al., 2019)), 25 repeats were run for each system. Thus, for 6 different C2 domains each with 3 different lipid membranes, a total of just under a millisecond of CG-MD simulations were performed. These were subsequently analysed in terms of the binding orientation of the C2 domain relative to the membrane, the dependence on the membrane lipid composition, and of the energetics of the interaction.

**Figure 2.**
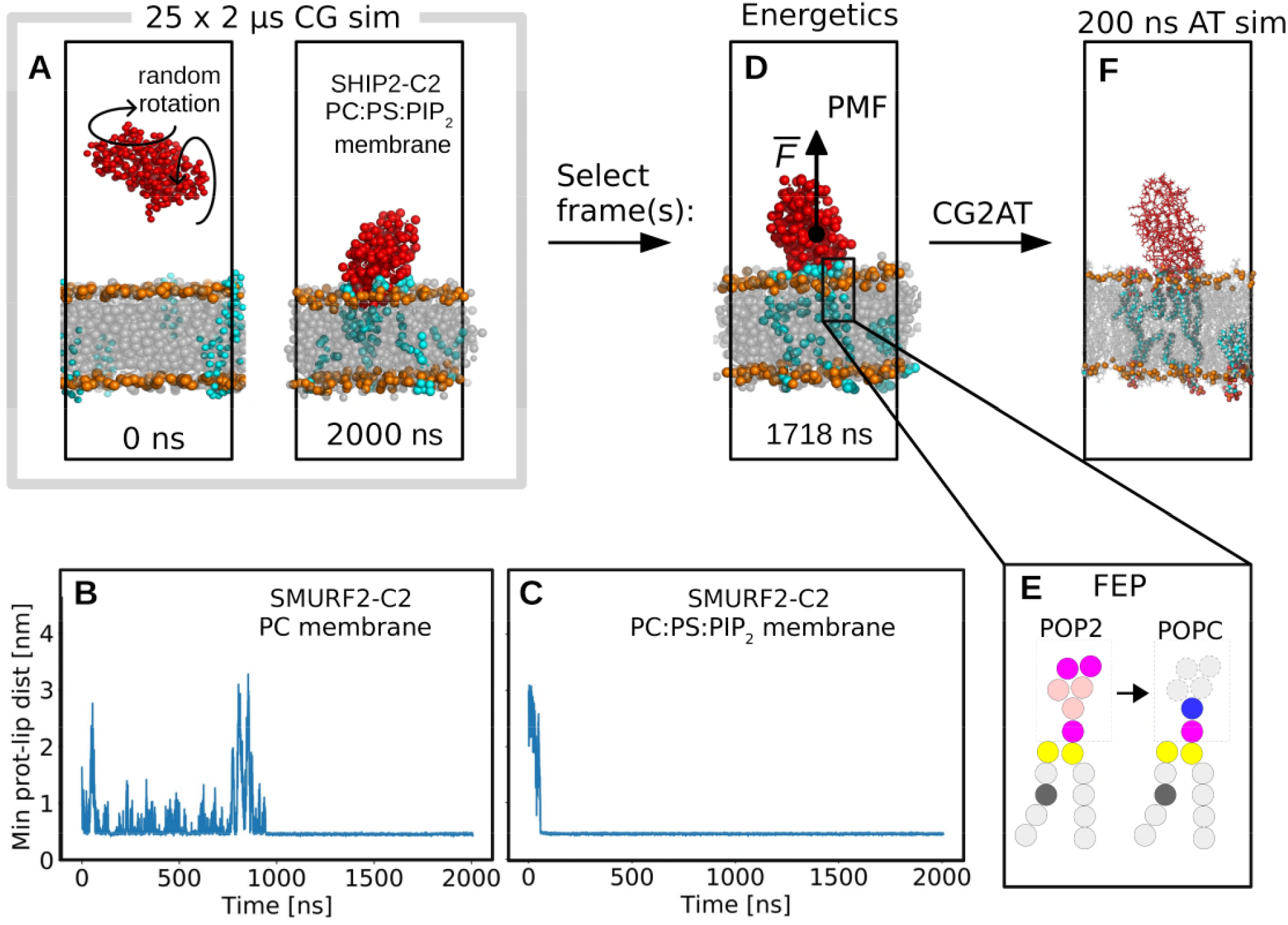
Simulation pipeline. **A** 25 replica CG simulations (each of 2 μs duration) of C2 domain/membrane association were run for each system. First and last frames are shown for the C2 from SHIP2 (red) with a PC:PS:PIP_2_ (80:15:5) membrane with PC and PS in grey, PIP_2_ in cyan, and beads representing phosphate groups in orange. **B, C** The minimum distance between the C2 domain protein and lipids of the bilayer for a single simulation (from an ensemble of 25 repeats) of the interaction of C2 from Smurf2 and either a PC membrane (**B**) or a PC:PS:PIP2 (80:15:5) membrane (**C**). The protein generally binds more quickly to the anionic membranes, and remains bound throughout the simulation, with some variation between replicas. **D** Representative binding mode(s) were selected for each system as described in the main text. Potentials of mean force (PMF), i.e. free energy profiles for the C2 domain/membrane interaction were calculated using umbrella sampling. **E** Free energy perturbation (FEP) calculations were performed in which PIP_2_ in the upper leaflet interacting with the bound C2 domain was converted to a PC molecule and the free energy change evaluated. **F** Representative binding modes were converted from CG (force field MARTINI 2.2) to atomistic (force field CHARMM 36m with TIP3P waters) representation and atomistic simulations were run for 200 ns.

### CG simulations of C2/membrane encounter

Six C2 domains were investigated, three type I and three type II (Table 1). The type I C2 domains were from KIBRA (PDB ID 6FJD), PI3KC2α (PDB ID 6BU0), and RIM2 (PDB ID 2BWQ), and the type II C2 domains from PTEN (PDB ID 1D5R), Smurf2 (PDB ID 2JQZ), and SHIP2 (PDB ID 5OKM). Here we will focus on a comparative overview of their simulated interactions with lipid bilayers of varying composition. Three types of membranes were included in the study: zwitterionic (phosphatidylcholine, PC), anionic (80% PC and 20% phosphatidylserine, PS), and anionic membrane including a PI lipids (80% PC, 15% PS, and 5% phosphatidylinositol bisphosphate, PIP_2_). We will use the shorthand notations PC, PC:PS, and PC:PS:PIP_2_ to denote these membranes. PC:PS:PIP_2_ is a model membrane for the negatively charged inner leaflet of mammalian plasma human membrane, which is relevant as all the C2 domains under consideration bind to this membrane *in vivo*.

**Table 1.**
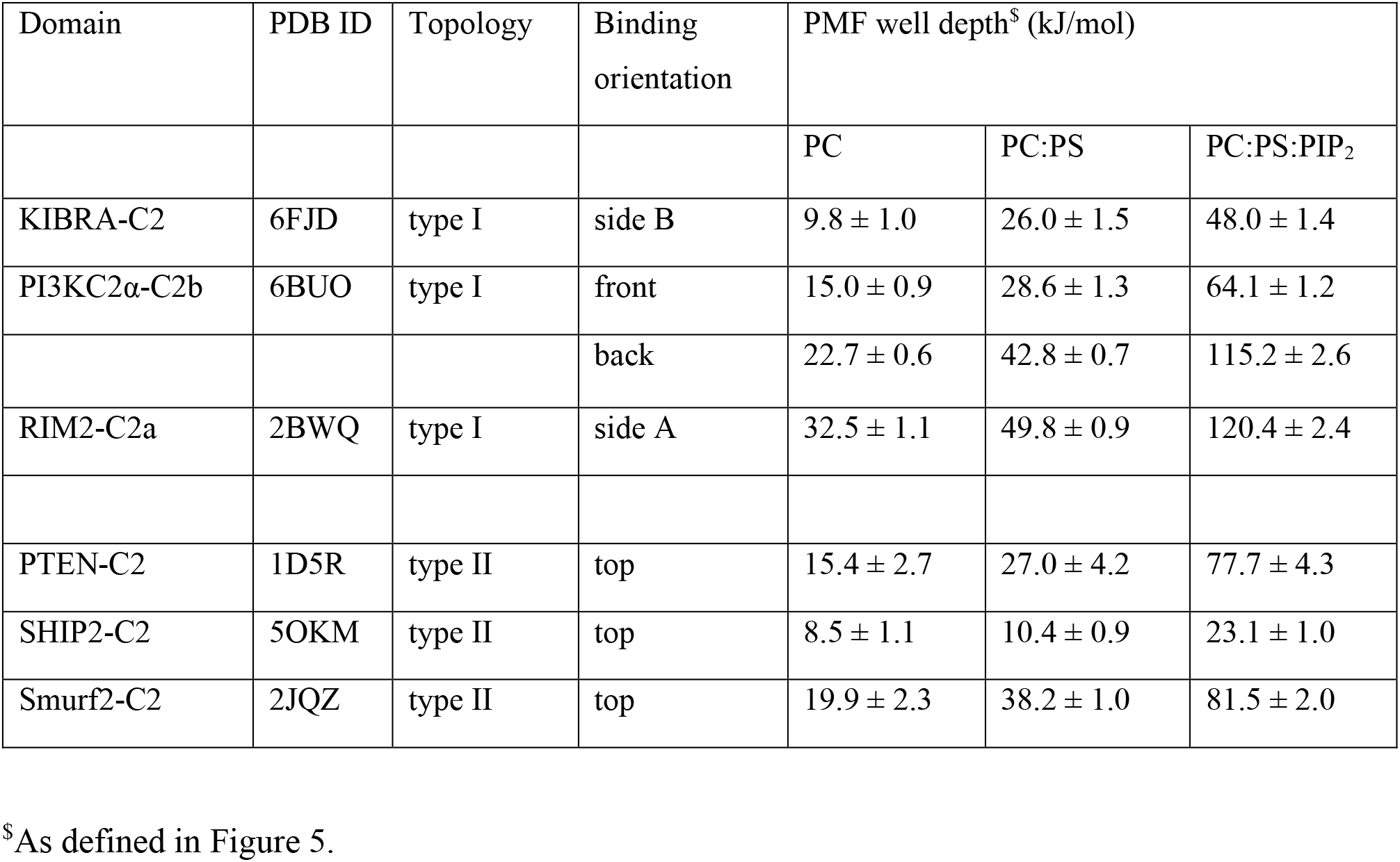
Summary of CG Association Simulations and PMFs.

As can be seen from the example of the Smurf2-C2 domain (Figure 2B-C) a single simulation in the presence of either a PC or a PC:PS:PIP_2_ bilayer can be used to demonstrate the influence of the lipid bilayer composition on the protein/membrane interaction, by monitoring the minimum distance between the protein and bilayer as a function of time. In this example, the C2 domain makes multiple encounters with the PC bilayer, before finally binding to the surface after ~1 μs (Figure 2B). In contrast, in the presence of the PI-containing bilayer, the initial encounter occurs within the first 0.1 μs (Figure 2C), leading to binding of the C2 domain to the membrane, with no subsequent dissociation over the course of the 2 μs simulation.

We repeated this analysis for the 25 repeats for all six C2 domains and 3 lipid compositions (Figure 3). Although the variation was relatively large between the repeats, especially for simulations with a PC bilayer, the overall pattern seen for the Smurf2-C2 is conserved. Thus, for all C2 domains the simulations with a PC bilayer lead to multiple reversible contacts between the C2 and the membrane. For the anionic PC:PS bilayer, in most cases multiple encounters are seen, on average leading to a longer lasting interaction of the protein and the membrane. In contrast, in the presence of a PC:PS:PIP_2_ bilayer, in all cases the initial encounter of C2 domain and bilayer leads to a protein/membrane interaction which persists for the remainder of the simulation. These simulations, especially those with PC:PS, enable us to come up with an initial, approximate ranking of the strength of interactions of C2 with a bilayer. Thus, comparing two C2 domains from PIP-phosphatases (i.e. PTEN and SHIP2) we may contrast PTEN-C2 which on average forms long-lasting interactions with both PC and PC:PS bilayers, with SHIP2-C2 which forms multiple reversible interactions with both PC and PC:PS bilayers and only forms long lasting interactions with a PC:PS:PIP2 bilayer. Fitting exponential decays to the averaged minimum protein-lipid distance vs. time data for these simulations (SI Figure S1 and Table S1) supports this interpretation.

**Figure 3.**
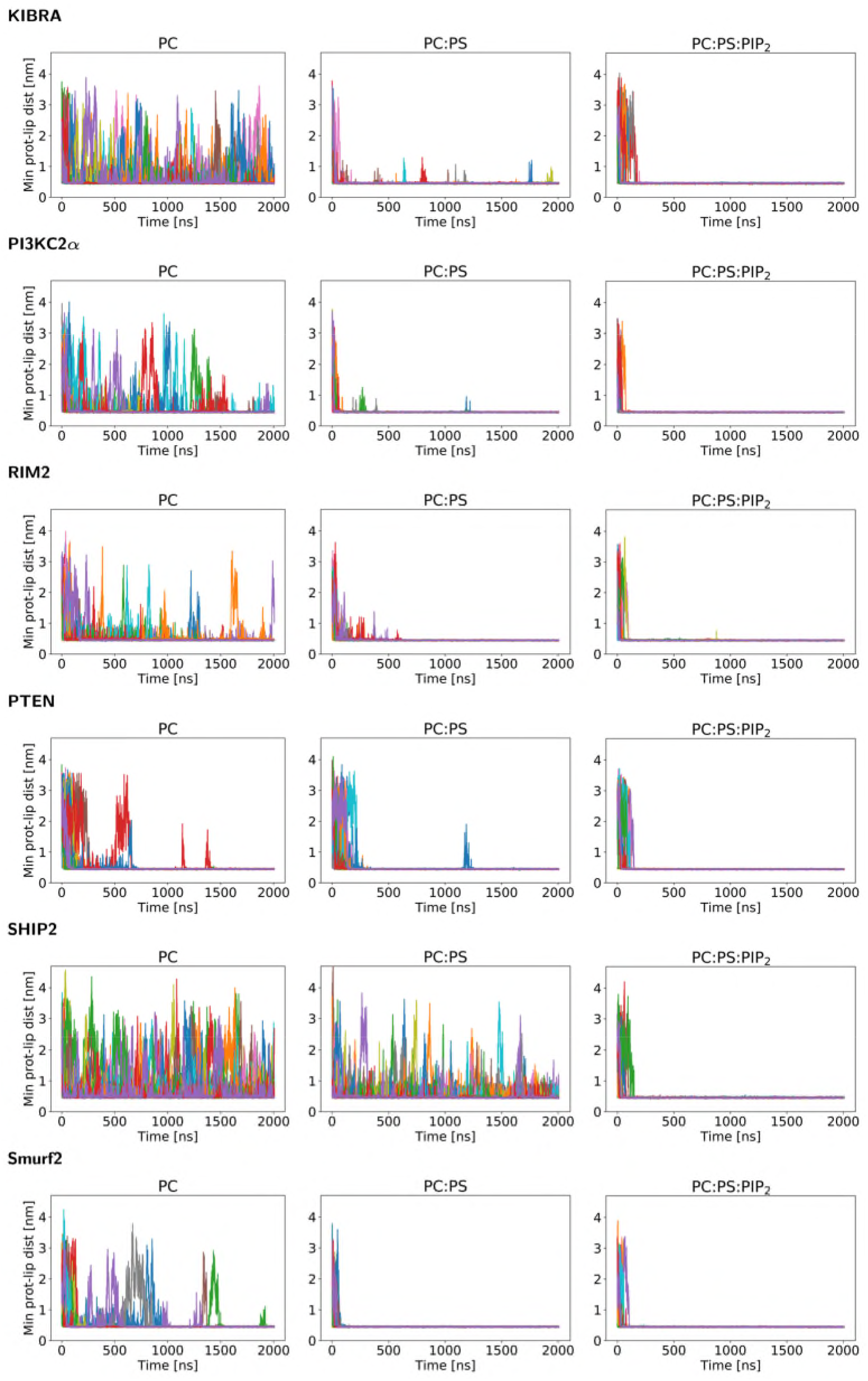
Minimum protein-lipid distances as a function of time for all simulations. For each simulation ensemble the minimum protein-lipid distance is shown as a function of time, with the different colours corresponding to the 25 repeats within the ensemble. Note that a distance of < 0.5 nm corresponds to a contract between the protein and the lipid bilayer. For each C2 domain, simulations are shown for interactions with a PC, a PC:PS, and a PC:PS:PIP_2_ bilayer.

The mode(s) of binding of the C2 domains were characterised by calculation of protein-lipid distance versus orientation density maps, which is an approach previously adopted for comparing PH domains (Yamamoto et al., 2016). The orientation of the C2 domain relative to the bilayer was quantified as the *zz* component of the rotation matrix, *R_zz_*, with respect to a reference frame (Figure 4). The last frame of the first repeat of the ensemble was used as a reference frame in the initial calculation of *R_zz_*, and subsequently all *R_zz_* values were recalculated with respect to a selected primary binding mode once this mode was determined. We found multiple binding modes for the isolated C2 domains, showing up as dense areas in the distance/orientation maps (Figure 4). In most cases, the modes were the same for zwitterionic membrane (PC), anionic membrane (PC:PS) and anionic membrane with PI (PC:PS:PIP_2_), and the most visited mode was in many cases the same for all membrane compositions (Figure 4). However, the C2 domains spend more time unbound and away from the membrane for the PC membrane, whilst the presence of PS and PI enhanced membrane-protein contacts.

**Figure 4.**
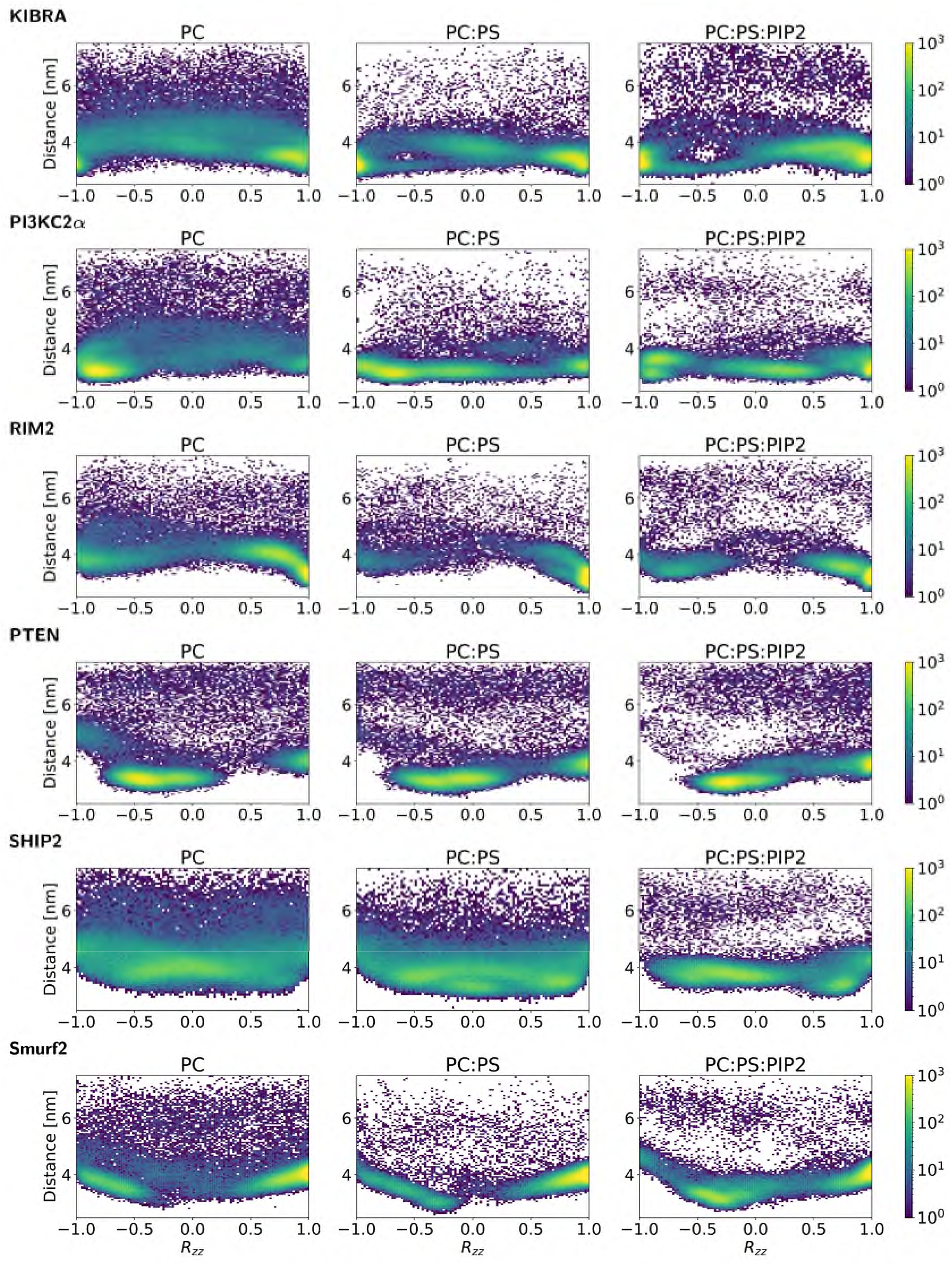
Density maps of the orientation and distance of the C2 domains relative to the lipid bilayer. For each simulation ensemble a density maps of the orientation and distance of the C2 domain relative to a lipid bilayer is given. Each density map represents the relative frequency (on a logarithmic colour scale from purple low to yellow high)), averaged across time and all 25 simulations in an ensemble, of the orientation and distance of the C2 domain relative to the bilayer. The orientation is given by *R_zz_* which is the *zz* component of the rotation matrix of the C2 domain with respect to a reference structure at *R_zz_* = 1 (see text for details of the reference structure). The distance shown is the *z*-component of the vector between the centres of mass of the lipid bilayer and of the C2 domain.

One primary binding mode was selected for each C2 domain (two for PI3KC2α). The selection criteria were: (i) the selected mode was *probable*, i.e. that the mode was frequently visited; (ii) the mode was *physically reasonable*, i.e. that domains adjacent to the C2 domain in the intact protein would not overlap with the membrane; (iii) the mode was *productive*, i.e. adjacent active domains were in contact with the bilayer. We note that criteria (ii) and (iii) could only be applied for C2 domains where the structure of adjacent domains was known.

### Two views of the energetics of C2/membrane-lipid interactions

To quantitatively compare the binding modes of the different C2 domains, we calculated binding free energies for each system. Firstly, a potential of mean force (PMF) was calculated using umbrella sampling with the centre-of-mass distance between protein and membrane as reaction coordinate (Figure 2D). This provides an estimate of a free energy curve for the interaction of the protein with the membrane. The C2 domains bound most strongly to PC:PS:PIP_2_ membrane, less tightly to anionic membrane without PIP_2_ (PC:PS), and even more loosely to PC membranes (Table 1 and Figure 5). The binding strengths had approximate ratios of 1:2:4 to 5 across the three membranes i.e. PC vs. PC:PS vs. PC:PS:PIP_2_ (Table 1).

**Figure 5.**
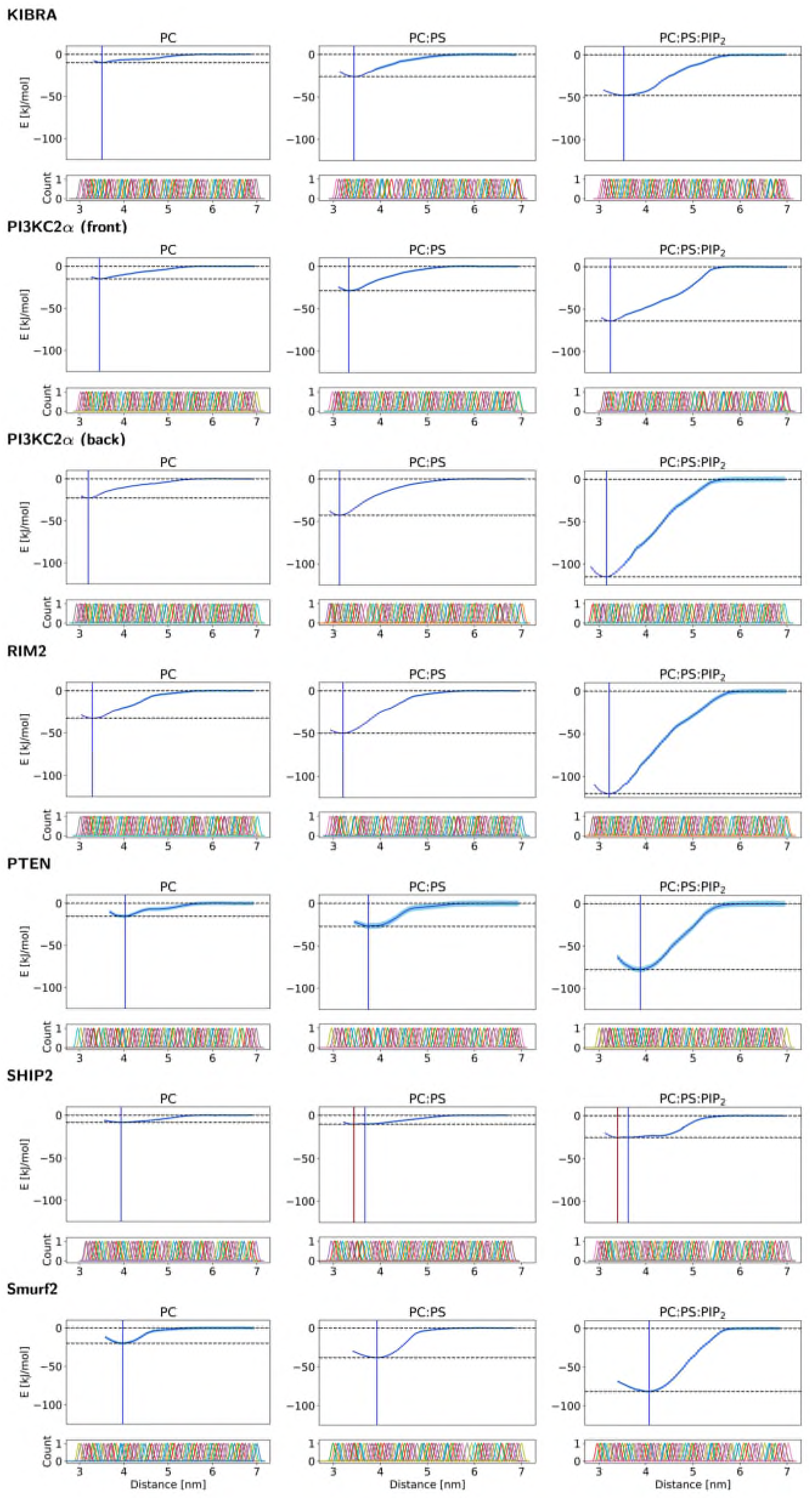
Potentials of mean force for C2 domain/membrane interactions. For each C2 domain and bilayer the potential of mean force (PMF) as calculated via umbrella sampling is shown, with the reaction coordinate corresponding to the distance between the centre of mass of the protein and centre of mass of the lipids. The dark blue curve shows the PMF with the pale blue are corresponding to ± one standard deviation. The two broken horizontal lines define the depth of the energy well corresponding to the minimum of the PMF. The corresponding histograms from the individual umbrella sampling window are shown beneath the PMF in order to demonstrate overlap.

Next, free energy perturbation (FEP) calculations provided additional information on the binding strength of specific PIP_2_ lipid head groups with the C2 domain (Figure 2E and Table 2). Four PIP_2_ molecules were bound to each C2 domain, with free energies ranging from −4 to −21 kJ/mol for each PIP_2_ molecule. Generally, the binding energies were evenly distributed among the four PIP_2_ headgroups bound in each system, with RIM2-C2 and the back-binding mode of PI3KC2α as notable exceptions (Table 2).

**Table 2:** Interaction free energies of PIP2 headgroups.

| Domain | Binding mode | FEP for each PIP<sub>2</sub> (kJ/mol)<sup>§</sup> | Total FEP (kJ/mol) | ΔPMF*<br>(kJ/mol) |
| --- | --- | --- | --- | --- |
| KIBRA-C2 | side B | 6, 7, 7, 6 | 26.0 ± 3.9 | 22.0 ± 2.1 |
| PI3KC2α-C2b | front | 9, 7, 10, 7 | 33.3 ± 4.0 | 35.5 ± 1.8 |
|  | back | 10, 7, 4, 10 | 30.9 ± 4.1 | 72.4 ± 2.7 |
| RIM2-C2a | side A | 11, 21, 15, 12 | 56.0 ± 3.6 | 70.6 ± 2.6 |
| PTEN-C2 | top | 10, 7, 9, 7 | 32.8 ± 4.0 | 50.7 ± 5.0 |
| SHIP2-C2 | top | 4, 4, 4, 4 | 15.7 ± 2.7 | 12.7 ± 1.3 |
| Smurf2-C2 | top | 6, 6, 9, 8 | 27.9 ± 4.4 | 43.3 ± 2.5 |

<sup>§</sup>Energy calculated by changing the headgroup of each PIP<sub>2</sub> in the upper leaflet to PC, whilst keeping the other PIP<sub>2</sub> molecules unaltered. Mean of 5 repeats. Errors on the total FEP is calculated via the standard deviation from the 5 repeats.
\* PMF well depth difference between binding to PC:PS (80:20) and binding to PC:PS:PIP<sub>2</sub> (80:15:5).

As an internal check, we both calculated binding energies via the potential of mean force (PMF) and free energy perturbation (FEP) (Figure 2D-E). Since all PIP_2_ in the upper leaflet were bound to C2, and these were converted to PC, the upper leaflet was effectively converted from a PC:PS:PIP_2_ (80:15:5) membrane to a PC:PS (85:15) membrane. Therefore, the FEP energies should be similar to the energy difference between the PMF for C2 bound to PC:PS:PIP_2_ (80:15:5) membranes and C2 bound to PC:PS (80:20) membranes (Table 2). The agreement is not perfect, but the trends are the same, and absolute values are close (Table 2). Binding energies were always smallest for binding to PC membranes, larger for PC:PS binding and largest for PC:PS:PIP_2_ membranes.

### Binding Modes, AT-MD, and comparison with experimental data

In the last step of the protocol, each system was converted from coarse-grain resolution to atomistic resolution, and a 200 ns simulation was performed (Figure 2F) to refine the binding modes (Figure 6). All C2 domains with type II topology bound with the top facing the membrane (Figure 6 and Table 1). The C2 domains with a type I topology, on the other hand, bound with either the side, front or back toward the membrane (Figure 1), i.e. with its longest axis parallel to the membrane. This is reflected in the PMF calculations through the distance with the minimum binding energy; the type I C2 domains have lowest energy at a center-of-mass distances between 3.1 and 3.5 nm, whereas the type II C2 domains have lowest energy at distances between 3.6 and 4.0 nm, for the productive mode (Figure 5). In all simulations with PIP_2_ present, the PIP_2_ molecules clustered around the C2 domain (Figure 6), with all four PIP_2_s in the upper leaflet bound for most of the time.

**Figure 6.**
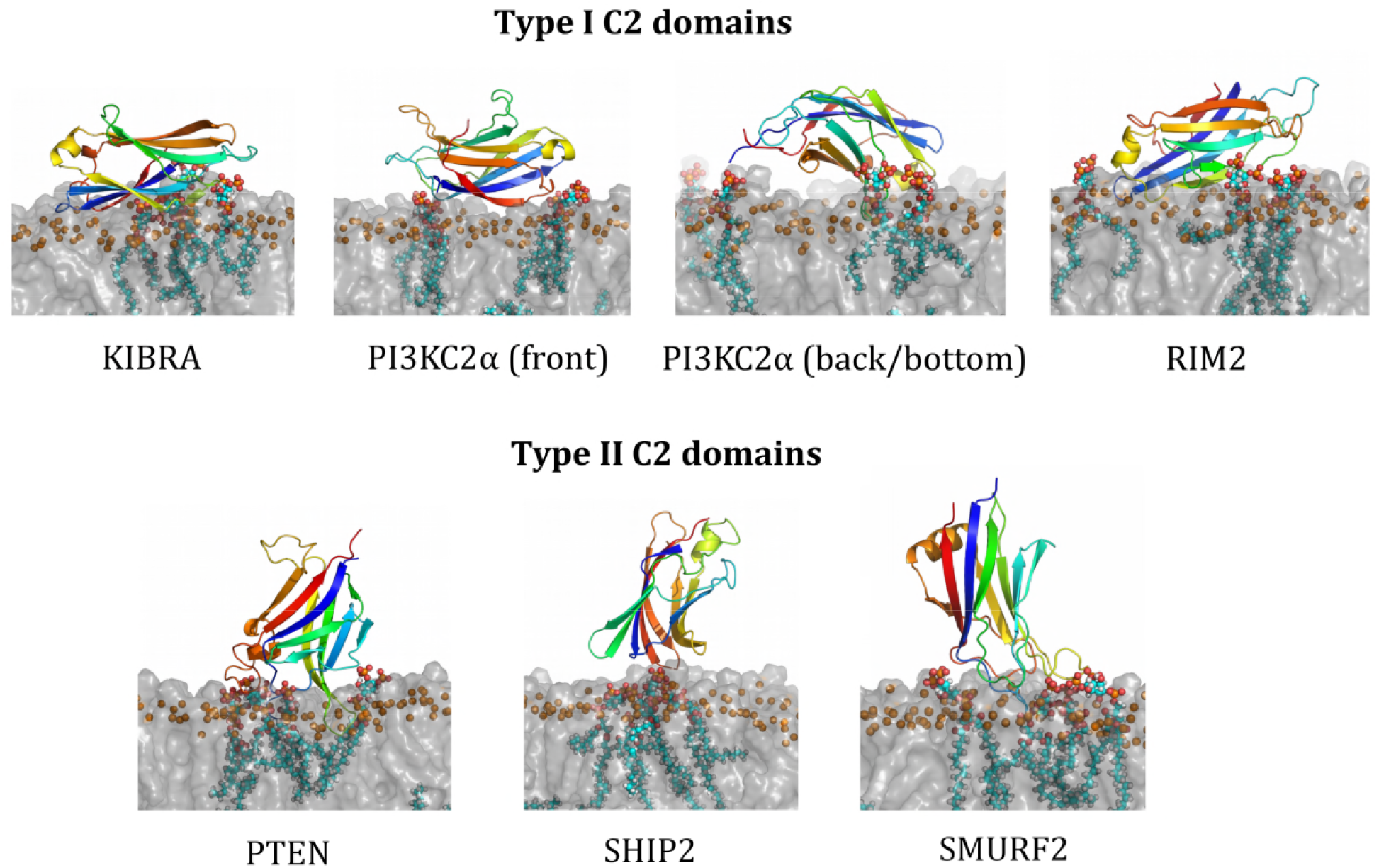
C2 membrane binding modes from atomistic simulations. Each panel shows a snapshot of the corresponding C2 domain at the end of an atomistic simulation whilst bound to a PC:PS:PIP_2_ bilayer, with POPC and POPS depicted as a transparent grey surface with their P atoms highlighted as orange spheres, and with PIP_2_ shows as van der Waals spheres. The C2 domains are shown as cartoon representations on a rainbow colour scheme from N-terminus (blue) to the C-terminus (red).

The C2 domains generally stayed in their initial bound configuration with PIP_2_ throughout the atomistic simulation, whereas C2 domains bound to PC:PS rotate more and unbind in one case, and C2 domains bound to pure PC membranes unbind in all but one case (Figure 7).

**Figure 7.**
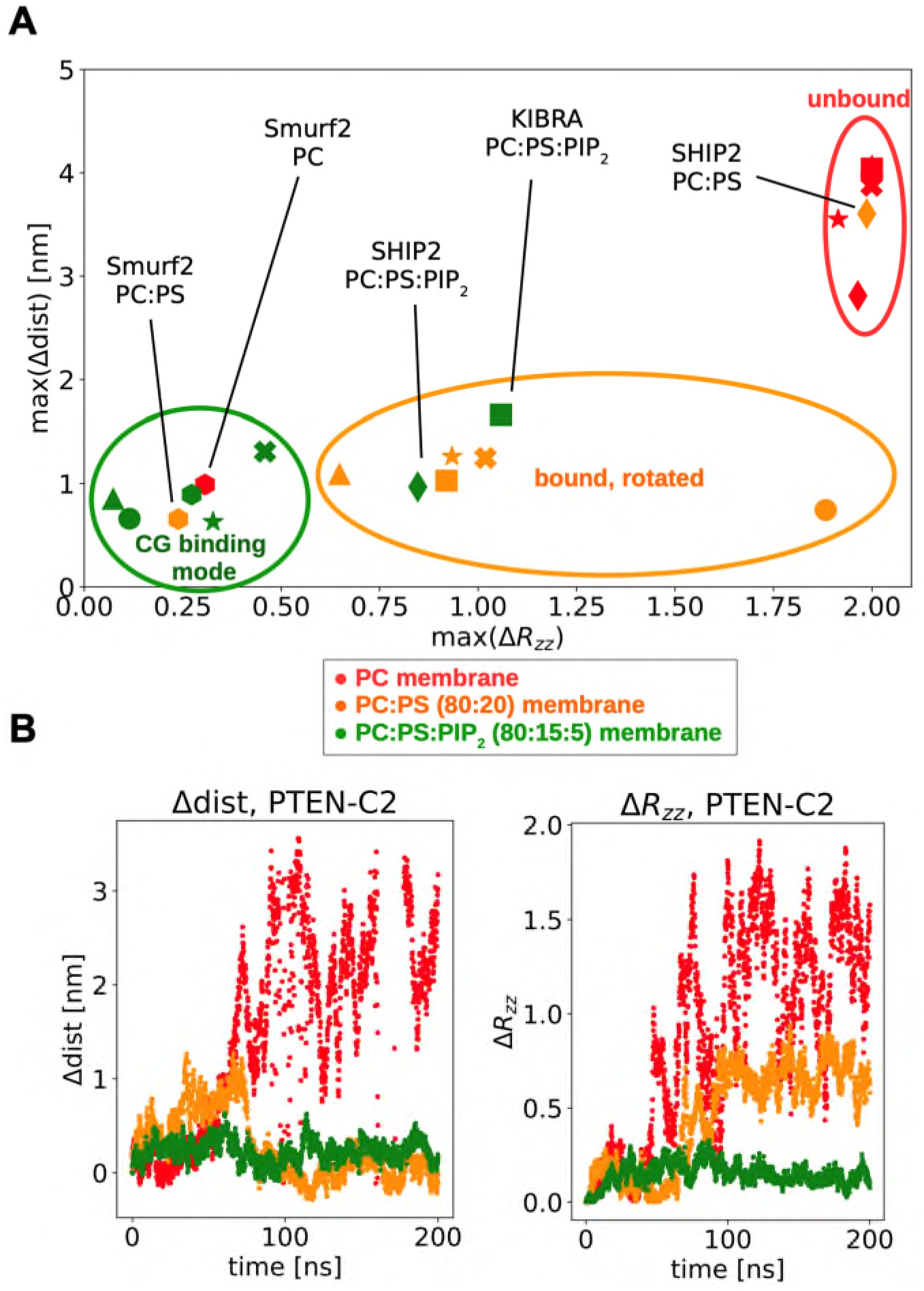
Changes in C2 orientation and distance relative the bilayer during atomistic simulations. **A** The maximum changes in distance and orientation (measured as *R_zz_* – see Methods) with respect to the initial frame (the CG binding mode) for the atomistic simulations. The simulations can be divided into three groups, shown by: a green ellipse, where the protein stay remains in the initial binding mode; an orange ellipse, where the protein remain bound but changes orientation (i.e. rotates relative to the bilayer); or a red ellipse, corresponding to those simulations where the C2 domain starts to dissociate from the membrane. Different point styles correspond to different C2 domains (KIBRA (square), PI3K (front mode, circle), PI3K (back mode, cross), RIM2 (triangle), PTEN (star), SHIP2 (diamond), and Smurf2 (hexagon)) and point colours to the lipids present in the bilayer (PC (red), PC:PS (orange), and PC:PS:PIP2 (green)). **B** An example of the change in distance and orientation during atomistic simulations of the PTEN-C2 domain initially bound to PC (red), PC:PS (orange) or PC:PS:PIP_2_ (green) bilayers.

In the following, we compare the binding modes observed in the simulations with structural and other experimental data where available.

#### Smurf2

The N-terminal C2 domain localizes Smurf (SMAD Specific E3 Ubiquitin Protein Ligase 2) to the membrane (Feng and Derynck, 2005; Kavsak et al., 2000; Wiesner et al., 2007). Smurf2-C2 showed one dominant binding mode in our CG simulations, which is the same for all three membranes (Figure 4). This is a canonical top-binding mode with loops L12, L56 and L78 facilitating lipid binding (Figure 6). These loops were previously determined as phospholipid binding sites by NMR (Wiesner et al., 2007), supporting our simulations. The Smurf-C2 stays in the same binding mode during 200 ns atomistic simulations for all three membranes (Figure 7). Notably, Smurf-C2 is the only protein out of the six investigated that stays bound to the PC membrane throughout the atomistic simulations. Smurf2-C2 shares 87% sequence identity with the C2 domain of Smurf1 (Figure S2), and they have similar functions and structure (Koganti et al., 2018), which make it relevant to compare them. Scott et al. used MD simulations to explore interactions of InsPs with Smurf1-C2 and they saw a canonical top-binding mode (Scott et al., 2020), in line with our results. Moreover, Smurf1-C2 has been crystallised with negatively charged sulphates bound, also in the top of the C2 domain (PDB ID 3PYC; SI Figure S2).

#### KIBRA-C2

Kidney and brain expressed protein (KIBRA), is a multi-functional protein with around 20 known binding partners and is central in many cellular processes, e.g. membrane trafficking (Milnik et al., 2012). KIBRA contains a C2 domain, which facilitates membrane binding in a Ca^2+^-independent manner (Posner et al., 2018). KIBRA-C2 displays two dominant binding modes in our simulations, but the most frequent binding mode is with β7 and β8 facing the bilayer (Figure 6), i.e. with side B (Figure 1) towards the lipid surface. This binding mode is consistent with previous NMR, X-ray crystallography and MD data for the lipid-binding site of KIBRA-C2 (Posner et al., 2018). The second binding mode (Figure 4) is unphysical, as the N- and C-termini are buried in the bilayer and KIBRA-C2 is a central domain. During the 200 ns atomistic simulation, the C2 domain changed orientation so the front faced the membrane (Figure 7) when bound to either PC:PS or PS:PS:PIP_2_ (the location of the front is shown in Figure 1). When initially bound to a PC membrane, the C2 domain diffused away from the bilayer.

#### RIM2-C2a

RIM2 (Rab3 interacting molecule 2) is involved in synaptic vesicle priming (Dai et al., 2005; Südhof, 2004). It contains two C2-domains, one in the middle of the protein (C2a) and one in the N-terminus (C2b). The structure of C2a has been solved and was studied here. RIM2-C2a has a dominant binding mode in our simulations with the front of the C2 domain binding the membrane (Figure 6) for all three membranes (Figure 4). A large binding energy to membranes for RIM2-C2 compared to the other C2 domains (Table 1) is likely due to a highly polar surface, with a positive patch on side A (β7 and β8) and a negative patch at side B (β4 and β5) (Dai et al., 2005), resulting in strong binding between side B and negatively charged membranes. RIM2 C2 has one particularly strongly bound PIP_2_ (with an FEP energy of 21 kJ/mol; Table 2). This lipid is bound to β4 of RIM2-C2, suggesting a specific PIP_2_ binding pocket at this location. In the atomistic simulations, the protein retained its orientation with a PIP_2_-containing membrane, underwent a rotation with PC:PS, and dissociated from the PC bilayer (Figure 7). RIM2-C2a has been crystallized with sulphates, that were suggested to bind at the same location as negatively charged PIP headgroups (PDB ID 2BWQ) (Dai et al., 2005). These sulphates are bound at a positively charged patch at the bottom of RIM2-C2a (SI Figure S3). In our simulations, a PIP_2_ headgroup was also bound at this site (SI Figure S3) (with binding energy of 11 kJ/mol; Table 2), suggesting that this is a specific PIP_2_ binding site.

#### PI3KC2α-C2b

PI3K (phosphoinositide-3-kinase) has a central C2-domain (C2a) and a C-terminal C2 domain (C2b). The structure of C2b has been solved (PDB ID 6BU0; note that C2b is referred to as PI3KC2_αC2C_ in (Chen et al., 2018)). PI2KC2α-C2b revealed two dominant binding modes in our simulations with PC:PS:PIP_2_ membranes, with, respectively, the front and the back of RIM-C2b facing the membrane (Figure 6). In the front binding mode, the lipids bind RIM-C2 mainly via the C-terminal β-strand, β8. In the back-binding mode, the lipids mainly bind with β4 and the loops L34 and L67, but also via β7 and L23. Both binding modes are possible, as they do not block the neighbouring PX domain in PI3KC2α, which is connected via a flexible loop at the N-terminus of C2 (Chen et al., 2018). The crystallographic asymmetric unit of 6BU0 contains three copies of the C2b domain and four InsP_6_ molecules which therefore define two distinct binding sites on the surface of the domain (Chen et al., 2018). The first binding site is at the front of the C2 domain, and the second is at the back, with L34 and L67 contacting the bound InsP_6_ molecule (SI Figure S4). Thus, the simulations and the crystal structure both suggest these two binding modes, i.e. a front- and a back-binding mode. The front-binding mode had four PIP_2_s bound in the simulations, but none of them coincided with the InsP_6_ from the crystal structure. Instead, the PIPs were bound close to the loops in the top and bottom of the C2 domain. The back-binding mode likewise had all four PIP_2_s from the upper leaflet bound, one of which was bound in proximity to the binding pocket of the crystal, i.e. the PIP_2_ headgroup from the simulation was bound about 0.8 nm from the crystallographic InsP_6_ (SI Figure S4). This suggests that this is may be a more specific binding pocket for PIP_2_, albeit having a relatively low binding affinity (4 kJ/mol; Table 2) as compared to the other bound PIPs.

#### PTEN-C2 and SHIP2-C2

Both of these C2 domains form the C-terminal half of a core enzyme structure, made up of a catalytic phosphatase (Ptase) domain followed by a membrane-targeting C2 domain (Le Coq et al., 2017; Worby and Dixon, 2014). This enables us to identify which binding modes of the isolated C2 domains are ‘productive’ (see above). Also, the simulated binding interactions of the isolated C2 domains may be compared with those of the core Ptase-C2 structures.

PTEN (phosphatase and tensin homolog) is a tumour suppressor which dephosphorylates PIP_3_ to PI(4,5)P_2_ (Worby and Dixon, 2014). The C2 domain of PTEN acts by bringing the phosphatase domain of PTEN (Ptase) close to the membrane. The isolated C2 from PTEN exhibits two binding modes in our simulations with all membranes (Figure 4). In the first mode, C2 binds with the loops in the top of the domain (Figure 6), whereas the second binding mode, which has the back of C2 interacting with the bilayer, is unphysical as the Ptase domain of PTEN would clash with the membrane (Lee et al., 1999) (PDB ID 1D5R). Therefore, the top binding mode is the only physical binding mode observed in our simulations.

SHIP2 (SH2-containing-inositol-5-phosphatase 2) dephosphorylates of PIP_3_ to PI(3,4)P_2_. The SHIP2-C2 exhibits three distinct binding-modes in simulations with a PC:PS:PIP_2_ membrane (Figure 4). The structure of SHIP-C2 has been determined with the adjacent Ptase domain (Le Coq et al., 2017) (PDB ID 5OKM), and in two of the modes, the Ptase domain is unproductively protruding out in the cytosol, with no contact to the lipids. In the productive mode, Ptase is close to the membrane, facilitated by C2. This latter binding mode corresponds to the top loops binding to the membrane, in particular L12, L34 and L56 (Figure 6). Interestingly, the PMF for SHIP2-C2 reveals that it binds weakly (Figure 5) with a broad minimum encompassing all 3 binding modes. Furthermore, even when bound to the PIP_2_-containing membrane, SHIP2-C2 has some rotational freedom in the atomistic simulation (Figure 7).

### Simulations of full-length proteins PTEN and SHIP2

As noted above, for PTEN and SHIP2 the structure of the core proteins (Ptase-C2) are known and so it is possible to also simulate these and compare the modes of membrane interactions with those of the isolated C2 domains. There have been a number of previous simulation studies of the interaction of PTEN with membranes (Gericke et al., 2013; Kalli et al., 2014; Lumb and Sansom, 2013; Nanda et al., 2015; S. Shenoy et al., 2012; S. S. Shenoy et al., 2012; Treece et al., 2020), but we have simulated it here again to maintain the same simulation settings as for isolated C2, in particular the same lipid composition, namely PC:PS:PIP_2_ (80:15:5).

As described above, for isolated PTEN-C2, two binding modes were observed, only one of which was ‘physical’, with the top loops pointing towards the bilayer. In the simulations of PTEN (Ptase+C2), we also got two binding modes (Figure 8A). One mode (~50% of the population; mode 1 in Figure 8A) was comparable to the ‘physical’ mode for the isolated C2 domain and allowed both the Ptase and C2 domains to contact the membrane (Figure 8A). An alternative mode 2 was also observed with side A of C2 pointing towards the lipids. This mode was not seen for the isolated C2. The catalytic site of Ptase of PTEN involves the P loop, the WDP loop and the TI loop (Lee et al., 1999), as well as an N-terminal motif covering residues 12-16 (NKRRY) (Gericke et al., 2013). In our simulated mode 1, the PIP_2_ molecules are in frequent contact with both the N-terminal motif, the P loop and the TI loop, and the PIPs have some contact to the WDP loop (Figure 8A). In the simulated mode 2 there is also contact between the PIPs and the N-terminal motif and the three loops, but substantially less (Figure 8A). So mode 1, which was also found for the isolated C2 domain, is likely to correspond to the active binding mode.

**Figure 8.**
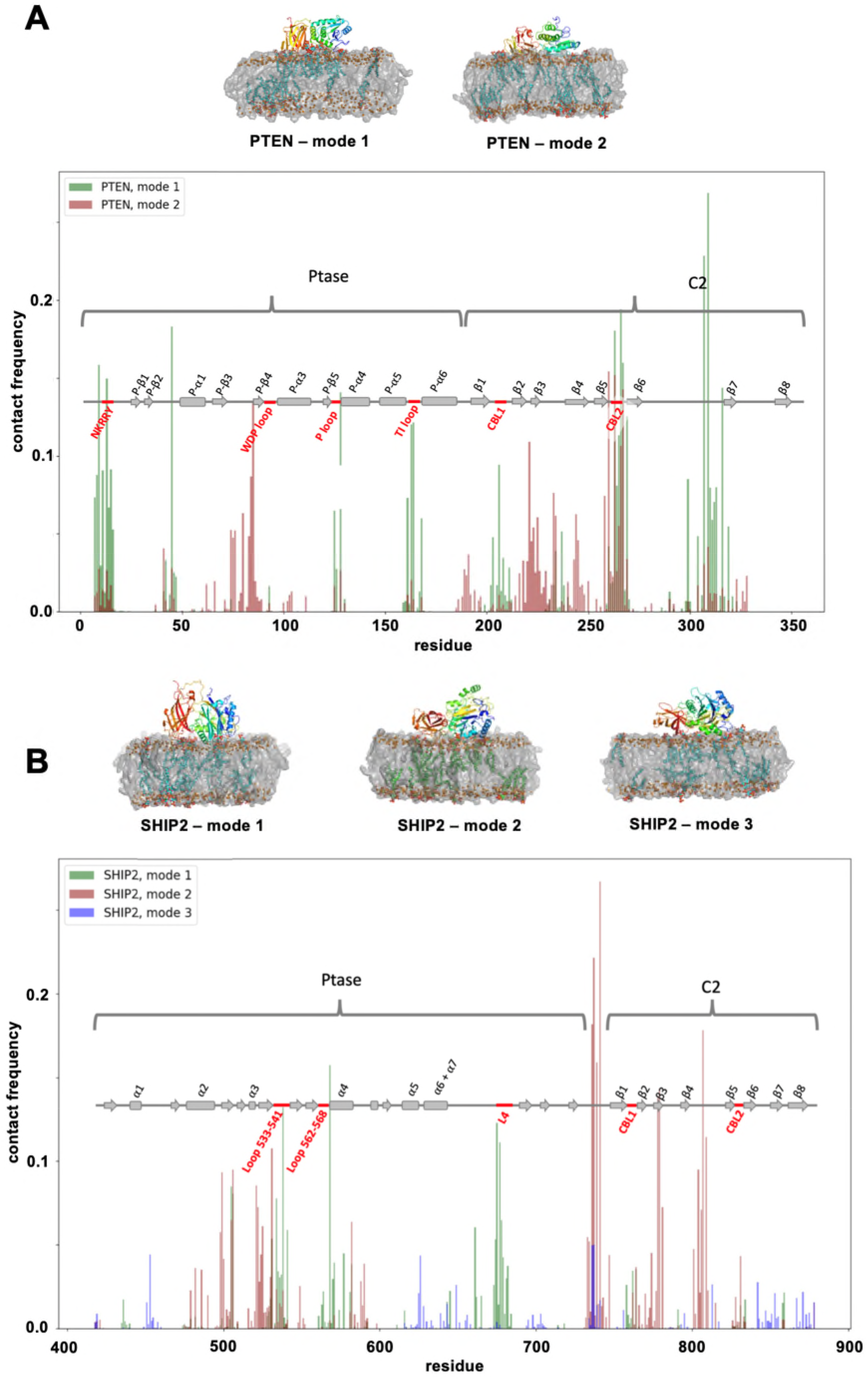
CG simulations of the interaction of intact (A) PTEN and (B) SHIP2 enzymes with PIP_2_-containing bilayers. In each case snapshots of the main modes of interaction are shown above graphs of the PIP_2_ contact frequencies as a function of residue number for the different binding modes. **A** Contact frequency (to PIP_2_) for each residue of PTEN. The two modes of PTEN binding are shown in green (mode 1) and red (mode 2). The secondary structure of PTEN is shown as: rectangles = α-helix and arrow = β-strand. The N-terminal domain, the WDP loop, the P loop and the TI loop (red) are all important for phosphatase activity. **B** Contact frequency for each residue of SHIP2. Three modes of SHIP2 binding are shown in green (mode 1), red (mode 2) and blue (mode 3). The loop L4, the loop containing residues 533-541 and the loop containing residues 562-568 (red) are part of the phosphatase active site.

For the isolated SHIP2-C2 domain, as discussed above, a rather weak multi-mode interaction was observed. For full-length SHIP2 (Ptase+C2) we observed three binding modes (Figure 8B). Of these, one mode (mode 1), similar to the productive mode for the isolated C2 accounted for 36 % of the population. In SHIP2, three loops are part of the catalytic site: the loop containing residues 675-684 (denoted ‘L4’ (Le Coq et al., 2017)), the loop consisting of residues 533-54,1 and the loop consisting of residues 562-568, as judged from available crystal structures (PDB IDs 5OKM and 4A9C) (Le Coq et al., 2017; Mills et al., 2012). Notably, in mode 1 of the simulations of full-length SHIP2, the Ptase has frequent contacts between the bound PIP_2_s and these three loops of the active site (Figure 8B). In the two other modes, on the other hand, there is little or no contact between PIP_2_s and the active site loops. Interestingly, in mode 1, the C2 seems to interact relatively weakly with the membrane, whereas the Ptase domain is more tightly bound, and PIP_2_ molecules cluster more around Ptase than around C2 (Figure 8B). This suggests that whilst the C2 domain may facilitate SHIP2/membrane interactions, in this case the Ptase domain is perhaps more important for the membrane interaction of SHIP2.

Comparing PTEN and SHIP2, these observations would suggest that whereas C2 from PTEN does, to some extent, determine the binding mode of PTEN, C2 from SHIP2 is less important in determining the binding orientation of SHIP2 and may just make a (weak) contribution to the avidity of binding. This is consistent with C2 from SHIP2 being the weakest binding C2 domain in the study (PMF = 23 kJ/mol), whereas C2 from PTEN has a higher membrane affinity (PMF = 78 kJ/mol).

## Discussion & Conclusions

We have demonstrated the utility of a simulation pipeline for exploring the interactions of Ca^2+^-independent C2 domains with anionic lipid bilayer models of cell membranes. This has allowed us to address a number of key questions, by comparing the interactions of six different species of C2 domain, namely: whether there is a systematic correlation between lipid binding and topology, how C2 domains bind to PIP molecules in a membrane, and whether C2 domains bind multiple PIP molecules, thus contributing to the formation of anionic lipid clusters. Here we review the implications of these findings in more detail.

### Non-canonical binding of C2 domains

C2 domains were first recognised as a Ca^2+^-dependent protein, with a canonical binding orientation involving the calcium binding loops at the “top” of the domain (Figure 1) (Corbalan-Garcia and Gómez-Fernández, 2014; Nalefski and Falke, 1996). Several studies have, however, expanded the possible modes to include binding without calcium and other studies have suggested that C2 domains can bind in a non-canonical orientation (Posner et al., 2018). We have systematically categorised and analysed some of these non-canonical binding modes. We found that all of the six domains studied bound to anionic membranes in the absence of Ca^2+^ ions, and that three of these proteins bound in non-canonical orientations via the front, back or side of the β-sandwich (Figures 6; consult Figure 1 for definitions of front, back, etc.). Notably, our results suggest a connection between topology and binding orientation, with type I C2 domains binding non-canonically, and type II C2 domains binding canonically, i.e. via the top loops.

### PIP_2_ binding and clustering

Clustering of PIP_2_ was observed on inspection of the final binding modes of the C2 domain (Figure 6). Binding of multiple PIPs to a single domain has also been reported for PH domains (Yamamoto et al., 2016), so this may be a general phenomenon for lipid-associated protein domains. Most of the binding energies for single PIP_2_ molecules are relatively small (<10 kJ/mol; Table 2), which may explain the lack of co-crystallised structures with e.g. inositol phosphates in the absence of the possibility to binding to multiple PIP_2_ in a membrane when in a crystal.

### Binding strength

We compared the binding strength of the C2 domains by different means. From the minimum distance vs. time data, we obtain the ranking (strongest binding first):

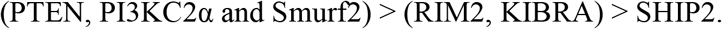

From the PMF calculations:

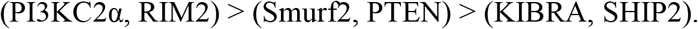

Finally, from the FEP calculations:

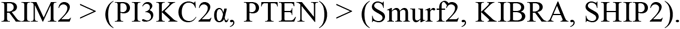

So, the consensus is: PI3KC2α, PTEN, RIM2 and Smurf2 form strong interactions whereas KIBRA, SHIP2 form weaker interactions. However, we note that (in part due to the limitations of the CG approach) we have not explored either PIP_2_ vs. PIP_3_ or PI(4,5)P_2_ vs. PI(3,4)P_2_ in terms of the strength of binding interactions. A more detailed examination of the strength and specificity of binding of PIP species would require development of robust atomistic simulation estimates of the energetics of protein binding to lipids (see (Pant and Tajkhorshid, 2020) for an example of this) and also a consideration of possible effects of ionization states of PIP headgroups and the influence of bound cations (see e.g. (Bilkova et al., 2017; Slochower et al., 2013)).

### Methodology

MD simulations allow for a systematic investigation of several related membrane recognition proteins, providing direct comparison between them. This would in most cases be challenging to do experimentally for six or more proteins. CG-MD allows us to monitor both binding of domains to bilayers of differing lipid composition and to undertake free energy calculations. The consistency observed for PMF and FEP calculations (Table 2) suggests that the energy calculations are converged. Convergence is an issue for energy calculation of protein membrane binding, as shown for PMF calculations of PH domains binding to a PIP-containing membrane (Naughton et al., 2018, 2016) and PIP_2_ bound to Kir2.2 integral membrane protein (Corey et al., 2019). In both cases, 500-1000 ns sampling was needed per umbrella window to obtain convergence (we used about 10 windows per protein/membrane combination in the present study). FEP needed 200-300 ns per window to obtain converge for PIP_2_ bound to Kir2.2 (Corey et al., 2019) (we used 20 windows). These accumulated simulation times are not feasible with AT-CG, so we did the energy calculations before converting to atomistic resolution. The CG force field may, on the other hand, not describe the energies accurately, so experimental benchmarking will be needed in the future to verify the absolute values of the energetics calculations.

We believe the established method is applicable to other membrane recognition domains. Similar computational approaches have already been exploited for comparative studies of PH domains (Naughton et al., 2016; Yamamoto et al., 2016), and the method can be applied to other domains as long as a sufficient number of solved structures are available.

There are hundreds of C2 domains (Nalefski and Falke, 1996), and the six included here are therefore not representative for all. We decided to limit the scope to include only C2 domains that (i) are Ca^2+^-independent, (ii) are interacting with the inner membrane leaflet, and (iii) are solved structurally. This limits the number of possible C2 domains to include. However, as new structures are solved, the methods can be applied to these, and results can be compared with the present results, to get more certain knowledge about the binding orientation, affinity and specificity of Ca^2+^-independent C2 domains.

### Conclusions

We have investigated six different Ca^2+^-independent C2 domains. They are all constituents of multidomain proteins that interact with the cell membrane and play a key role in signalling. Using multi-scale molecular dynamics, we investigated their binding modes, including binding orientation, binding affinity, and lipid specificity. We found that binding orientation and structural topology were related: type I C2 domains bound to the anionic membranes via the sides, front or back of the β-sandwich, whereas type II C2 domains bound via the loops at the top of the structure. Calculated binding free energies revealed significant binding differences between the six C2 domains, but these binding energies were not systematically correlated with the domain topology. Moreover, the domains, in general, only bound and stayed bound to negatively charged membranes, i.e. those containing PS and/or PIP_2_, and bound significantly more strongly to PIP_2_-containing membranes. PIP_2_ clustered around the C2 domains upon binding, which suggests that C2 lipid-interaction generally involves binding of several clustered PIPs, as also observed for the related PH domain (Yamamoto et al., 2016). PS, on the other hand, did not cluster around the C2-domains, nor did PC. We have thus provided an overview of expected binding modes, binding affinity and binding specificity for non-canonical C2-domains, which improves our overall understand the cellular roles of C2 domains.

Equally important, we have presented a transferable methodology that can be utilised for novel structures of C2 domains and can readily be expanded for investigations of other protein domains interacting with lipid bilayer models of cell membranes. Advances in both spatially resolved lipidomics (Tsuji et al., 2019) and in simulations of complex membrane models (Marrink et al., 2019) combined with systematic simulation methodologies will in the future allow us to reliably predict patterns of protein/membrane recognition (Irvine et al., 2019).

## Methods

### Coarse-grained molecular dynamics (CG-MD) simulations

CG-MD simulations were done using the Martini 2.2 force field (de Jong et al., 2013) and performed in GROMACS 2018.6 (Abraham et al., 2015). The input structures were truncated to only contain the C2 domain, using PyMOL (The PyMOL Molecular Graphics System, version 2.0 Schrödinger, LLC) and missing residues were added using Modeller (Fiser et al., 2000). The protein was coarse-grained using the Martinize script (de Jong et al., 2013) with the default elastic network to maintain the tertiary structure. The protein was positioned above a lipid membrane using the Insane script (Wassenaar et al., 2015) with box size 7×7×18 nm^3^. The protein placed at a minimum distance of 4.4 nm from the membrane, which is four times the VDW and electrostatic cutoff distances of 1.1 Å. 10% antifreeze water was added (Marrink et al., 2007) and the system was neutralized with Na^+^ or Cl^−^ ions. The protein was rotated through a randomly selected angle relative to the bilayer to avoid any bias in biding orientation. Three different membranes were generated for each protein: PC, PC:PS (molar ratio 80:20) and PC:PS:PIP_2_ (80:15:5). For our box size, 5% PIP_2_ corresponds to four molecules in each leaflet. We used the Martini lipid POP2 for PIP_2_ (López et al., 2013). The system was first minimised, then minimized using a restraint to ensure that the protein could only encounter the upper leaflet of the membrane. The restraint was set up using PLUMED UPPER_WALLS (Tribello et al., 2014) at a protein-lipid centre-of-mass distance of 7 nm and an energy constant of κ = 50 (internal units of code). Equilibration was run in the NPT ensemble with a semi-isotropic Berendsen barostat (Berendsen et al., 1984) and time constant of 14 ps to keep pressure at 1 bar, and at v-rescale temperature coupling applied separately to lipids, protein and solvent, with a time constant of 1 ps to keep the temperature at 323 K. Equilibration was run with 20 fs time steps for 10 ns, and production runs were run with the same settings, but with 35 fs time steps and for 2 μs. 25 repeats were made for each protein/membrane combination, with a new rotation, and setup of the system for each repeat.

### Generating distance vs. R_zz_ plots

Distances between the centre of mass of the protein and the centre of mass of the lipids were calculated for each frame using *gmx distance* (after centering the protein in the box). A reference frame was needed for calculating the rotation matrix. One frame for each protein was selected from one of the 25 simulations with membrane PC:PS:PIP_2_. The protein from this frame was extracted, and used as reference protein, also for simulations with PC and PC:PS membranes. The rotation matrix was then calculated for each frame by comparing the protein in each frame with the reference protein using *gmx rotmat*, after fitting in the *xy* plane. R_zz_ is the zz component of the rotation matrix.

### Potential of Mean Force (PMF) calculations

PMFs were calculated using umbrella sampling with the centre-of-mass distance between protein and membrane as reaction coordinate (Figure 2E). As preparation for the PMF calculations, two steered MD simulation was performed to generate starting frames for the umbrella sampling along the reaction coordinate, using the GROMACS pull code (GROMACS, 2020). In the first, the protein was pushed into the bilayer, and in the second, the protein was pulled away from the bilayer. The distance between the centre of mass of the protein and centre of mass of the lipids was restrained with a harmonic potential with force constant of 1000 kJ/mol/nm^2^ and a rate of 0.2 nm/ns in the *z*-direction (positive and negative direction for push and pull simulation, respectively). A position restraint with a force constant of 15,000 kJ/mol/nm^2^ was applied on the strongly bound PIP_2_ lipids to prevent them from being pulled out of the membrane with the protein. Frames were retrieved from the steered MD run every 0.05 nm until a centre-of-mass distance of 7.0 nm was obtained for the last frame (about 10 windows per system). Umbrella sampling was then performed by sampling each retrieved frames for 1 μs with a 2000 kJ/mol/nm^2^ position restraint on the protein, but with the restraint on PIP_2_ removed. The PMF was calculated using *gmx wham* (Hub et al., 2010), skipping the first 200 ns as equilibration, and using the bootstrap method (option *nBootstrap*) to get uncertainties on the PMF values.

### Free Energy Perturbations (FEP)

Free energy perturbations were done as previously described (Corey et al., 2020, 2019), converting the Martini lipid POP2 to Marini lipid POPC (Figure 2F). The head group beads were gradually changed, in 20 steps, as controlled by a parameter λ. Each of the four POP2 Martini lipids in the upper leaflet was converted individually to POPC, with the others kept as POP2. To keep the system neutral at all times, we gradually converted five sodium ions to neutral beads. Each frame was sampled for 1 μs, and the free energy was calculated using the *alchemical_analysis* script and the MBAR method (Klimovich et al., 2015). The perturbation energies were also calculated for conversion of POP2 in a bilayer without bound protein. The reported energies are the difference between FEP energies for free and bound POP2 (Corey et al., 2019).

### Atomistic molecular dynamics (AT-MD) simulations

Selected frames with a membrane-bound C2 molecule were converted from CG (force field Martini 2.2) to AT (force field CHARMM36m (Best et al., 2012; Huang et al., 2017) and TIP3P water) using the CG2AT script (Vickery and Stansfeld, 2020). Atomistic simulations were run in GROMACS 2018.6 (Abraham et al., 2015). The system was first minimized, then equilibrated for 100 ps in the NVT ensemble and 100 ps in the NPT ensemble. The equilibrated system was run for 200 ns. Both equilibration steps and production run had 2 fs timesteps with 1.2 nm VDW and electrostatic cutoff distance, v-rescale temperature coupling keeping the temperature at 300 K with timeconstant of ps, separately for protein, lipids and solvent. A Parrinello-Rahman barostat (Parrinello and Rahman, 1981) was applied to the NPT equilibration and production run with a semiisotropic pressure coupling, keeping pressure at 1 bar with time constant of 5 ps and water compressibility of 4.5×10^−5^ bar^−1^.

## Acknowledgements

The authors would like to thank Robin Corey for input on running free energy calculations, Owen Vickery for extending the CG2AT script, and Sarah-Beth Amos and Laura John for providing useful scripts for the analysis. AHL was funded by the Carlsberg foundation (CF19-0288). The authors acknowledge Wellcome (208361/Z/17/Z), BBSRC (BB/R00126X/1) and HECBioSim (EPSRC, EP/R029407/1) for funding.

## References

Abraham MJ, Murtola T, Schulz R, Páll SS, Smith JC, Hess B, Lindahl E, Lindah E. 2015. Gromacs: High performance molecular simulations through multi-level parallelism from laptops to supercomputers. SoftwareX 1–2:19–25. doi:10.1016/j.softx.2015.06.001

Alwarawrah M, Wereszczynski J. 2017. Investigation of the Effect of Bilayer Composition on PKcα-C2 Domain Docking Using Molecular Dynamics Simulations. J Phys Chem B 121:78–88. doi:10.1021/acs.jpcb.6b10188

Amos SBTA, Kalli AC, Shi J, Sansom MSP. 2019. Membrane Recognition and Binding by the Phosphatidylinositol Phosphate Kinase PIP5K1A: A Multiscale Simulation Study. Structure 27:1336–1346.e2. doi:10.1016/j.str.2019.05.004

Banci L, Cavallaro G, Kheifets V, Mochly-Rosen D. 2002. Molecular dynamics characterization of the C2 domain of protein kinase Cβ*. J Biol Chem 277:12988–12997. doi:10.1074/jbc.M106875200

Berendsen HJC, Postma JPM, Van Gunsteren WF, Dinola A, Haak JR. 1984. Molecular dynamics with coupling to an external bath. J Chem Phys 81:3684–3690. doi:10.1063/1.448118

Best RB, Zhu X, Shim J, Lopes PEM, Mittal J, Feig M, MacKerell AD. 2012. Optimization of the additive CHARMM all-atom protein force field targeting improved sampling of the backbone φ, ψ and side-chain χ1 and χ2 Dihedral Angles. J Chem Theory Comput 8:3257–3273. doi:10.1021/ct300400x

Bilkova E, Pleskot R, Rissanen S, Sun S, Czogalla A, Cwiklik L, Róg T, Vattulainen I, Cremer PS, Jungwirth P, Coskun Ü. 2017. Calcium Directly Regulates Phosphatidylinositol 4,5-Bisphosphate Headgroup Conformation and Recognition. J Am Chem Soc 139:4019–4024. doi:10.1021/jacs.6b11760

Chen KE, Tillu VA, Chandra M, Collins BM. 2018. Molecular Basis for Membrane Recruitment by the PX and C2 Domains of Class II Phosphoinositide 3-Kinase-C2α. Structure 26:1612–1625. doi:10.1016/j.str.2018.08.010

Chon NL, Osterberg JR, Henderson J, Khan HM, Reuter N, Knight JD, Lin H. 2015. Membrane Docking of the Synaptotagmin 7 C2A Domain: Computation Reveals Interplay between Electrostatic and Hydrophobic Contributions. Biochemistry 54:5696–5711. doi:10.1021/acs.biochem.5b00422

Corbalan-Garcia S, Gómez-Fernández JC. 2014. Signaling through C2 domains: More than one lipid target. Biochim Biophys Acta - Biomembr 1838:1536–1547. doi:10.1016/j.bbamem.2014.01.008

Corey RA, Stansfeld PJ, Sansom MSP. 2020. The energetics of protein-lipid interactions as viewed by molecular simulations. Biochem Soc Trans 48:25–37. doi:10.1042/BST20190149

Corey RA, Vickery ON, Sansom MSP, Stansfeld PJ. 2019. Insights into Membrane Protein-Lipid Interactions from Free Energy Calculations. J Chem Theory Comput 15:5727–5736. doi:10.1021/acs.jctc.9b00548

Corradi V, Sejdiu BI, Mesa-Galloso H, Abdizadeh H, Noskov SY, Marrink SJ, Tieleman DP. 2019. Emerging Diversity in Lipid-Protein Interactions. Chem Rev 119:5775–5848. doi:10.1021/acs.chemrev.8b00451

Dai H, Tomchick DR, García J, Südhof TC, Machius M, Rizo J. 2005. Crystal structure of the RIM2 C2A-domain at 1.4 Å resolution. Biochemistry 44:13533–13542. doi:10.1021/bi0513608

de Jong DH, Singh G, Bennett WFD, Arnarez C, Wassenaar TA, Schäffer L V, Periole X, Tieleman DP, Marrink SJ. 2013. Improved Parameters for the Martini Coarse-Grained Protein Force Field. J Chem Theory Comput 9:687–697. doi:10.1021/ct300646g

Feng XH, Derynck R. 2005. Specificity and versatility in TGF-β signaling through smads. Annu Rev Cell Dev Biol 21:659–693. doi:10.1146/annurev.cellbio.21.022404.142018

Fiser A, Do RK, Sali A, Fiser András, Kinh R, Do G, Andrej S. 2000. Modeling of loops in protein structures. Protein Sci 9:1753–1773. doi:10.1110/ps.9.9.1753

Gericke A, Leslie NR, Lösche M, Ross AH. 2013. PtdIns(4,5)P2-Mediated Cell Signaling: Emerging Principles and PTEN as a Paradigm for Regulatory Mechanism In: Capelluto DGS, editor. Lipid-Mediated Protein Signaling. Dordrecht: Springer Netherlands. pp. 85–104. doi:10.1007/978-94-007-6331-9_6

GROMACS. 2020. GROMACS pull code. http://manual.gromacs.org/documentation/current/user-guide/mdp-options.html

Hedger G, Sansom MSP. 2016. Lipid interaction sites on channels, transporters and receptors: Recent insights from molecular dynamics simulations. Biochim Biophys Acta - Biomembr 1858:2390–2400. doi:10.1016/j.bbamem.2016.02.037

Huang J, Rauscher S, Nawrocki G, Ran T, Feig M, de Groot BL, Grubmüller H, MacKerell AD. 2017. CHARMM36m: An Improved Force Field for Folded and Intrinsically Disordered Proteins. Nat Methods 14:71–73. doi:10.1038/nmeth.4067

Hub JS, De Groot BL, Van Der Spoel D. 2010. g_whams - A Free Weighted Histogram Analysis Implementation Includin Robust Error and Autocorrelation Estimates. J Chem Theory Comput 6:3713–3720. doi:10.1021/ct100494z

Ingólfsson HI, Lopez CA, Uusitalo JJ, de Jong DH, Gopal SM, Periole X, Marrink SJ. 2014. The power of coarse graining in biomolecular simulations. Wiley Interdiscip Rev Comput Mol Sci 4:225–248. doi:10.1002/wcms.1169

Irvine WA, Flanagan JU, Allison JR. 2019. Computational Prediction of Amino Acids Governing Protein-Membrane Interaction for the PIP3 Cell Signaling System. Structure 27:371–380. doi:10.1016/j.str.2018.10.014

Jaud S, Tobias DJ, Falke JJ, White SH. 2007. Self-induced docking site of a deeply embedded peripheral membrane protein. Biophys J 92:517–524. doi:10.1529/biophysj.106.090704

Kalli AC, Devaney I, Sansom MSP. 2014. Interactions of phosphatase and tensin homologue (PTEN) proteins with phosphatidylinositol phosphates: Insights from molecular dynamics simulations of PTEN and voltage sensitive phosphatase. Biochemistry 53:1724–1732. doi:10.1021/bi5000299

Kalli AC, Sansom MSP. 2014. Interactions of peripheral proteins with model membranes as viewed by molecular dynamics simulations. Biochem Soc Trans 42:1418–1424. doi:10.1042/BST20140144

Katan M, Allen VL. 1999. Modular PH and C2 domains in membrane attachment and other functions. FEBS Lett 452:36–40. doi:10.1016/S0014-5793(99)00531-1

Kavsak P, Rasmussen RK, Causing CG, Bonni S, Zhu H, Thomsen GH, Wrana JL. 2000. Smad7 Binds to Smurf2 to Form an E3 Ubiquitin Ligase that Targets the TGFbeta Receptor for Degradation. Mol Cell 6:1365–1375.

Klimovich P V., Shirts MR, Mobley DL. 2015. Guidelines for the analysis of free energy calculations. J Comput Aided Mol Des 29:397–411. doi:10.1007/s10822-015-9840-9

Koganti P, Levy-Cohen G, Blank M. 2018. Smurfs in protein homeostasis, signaling, and cancer. Front Oncol 8:295. doi:10.3389/fonc.2018.00295

Kutateladze TG. 2010. Translation of the phosphoinositide code by PI effectors. Nat Chem Biol 6:507–513. doi:10.1038/nchembio.390

Lai CL, Landgraf KE, Voth GA, Falke JJ. 2010. Membrane docking geometry and target lipid stoichiometry of membrane-bound PKCα C2 domain: A combined molecular dynamics and experimental study. J Mol Biol 402:301–310. doi:10.1016/j.jmb.2010.07.037

Lai CL, Srivastava A, Pilling C, Chase AR, Falke JJ, Voth GA. 2013. Molecular mechanism of membrane binding of the GRP1 PH domain. J Mol Biol 425:3073–3090. doi:10.1016/j.jmb.2013.05.026

Le Coq J, Camacho-Artacho M, Velázquez J, Santiveri CM, Gallego LH, Campos-Olivas R, Dölker N, Lietha D. 2017. Structural basis for interdomain communication in SHIP2 providing high phosphatase activity. Elife 6:1–25. doi:10.7554/eLife.26640

Lee JO, Yang H, Georgescu MM, Cristofano A Di, Maehama T, Shi Y, Dixon JE, Pandolfi P, Pavletich NP. 1999. Crystal structure of the PTEN tumor suppressor: Implications for its phosphoinositide phosphatase activity and membrane association. Cell 99:323–334. doi:10.1016/S0092-8674(00)81663-3

López CA, Sovova Z, Van Eerden FJ, De Vries AH, Marrink SJ. 2013. Martini force field parameters for glycolipids. J Chem Theory Comput 9:1694–1708. doi:10.1021/ct3009655

Lumb CN, Sansom MSP. 2013. Defining the membrane-associated state of the PTEN tumor suppressor protein. Biophys J 104:613–621. doi:10.1016/j.bpj.2012.12.002

Maehama T, Dixon JE. 1999. PTEN: A tumour suppressor that functions as a phospholipid phosphatase. Trends Cell Biol 9:125–128. doi:10.1016/S0962-8924(99)01519-6

Manna D, Bhardwaj N, Vora MS, Stahelin R V., Lu H, Cho W. 2008. Differential roles of phosphatidylserine, PtdIns(4,5)P2, and PtdIns(3,4,5)P3 in plasma membrane targeting of C2 domains: Molecular dynamics simulation, membrane binding, and cell translocation studies of the PKCα C2 domain. J Biol Chem 283:26047–26058. doi:10.1074/jbc.M802617200

Manna M, Nieminen T, Vattulainen I. 2019. Understanding the Role of Lipids in Signaling Through Atomistic and Multiscale Simulations of Cell Membranes. Annu Rev Biophys 48:421–439. doi:10.1146/annurev-biophys-052118-115553

Marrink SJ, Corradi V, Souza PCT, Ingólfsson HI, Tieleman DP, Sansom MSP. 2019. Computational Modeling of Realistic Cell Membranes. Chem Rev 119:6184–6226. doi:10.1021/acs.chemrev.8b00460

Marrink SJ, Risselada HJ, Yefimov S, Tieleman DP, de Vries AH. 2007. The MARTINI Force Field: Coarse Grained Model for Biomolecular Simulations. J Phys Chem B 111:7812–7824.

Marrink SJ, Tieleman DP. 2013. Perspective on the Martini model. Chem Soc Rev 42:6801–6822. doi:10.1039/c3cs60093a

Michaeli L, Gottfried I, Bykhovskaia M, Ashery U. 2017. Phosphatidylinositol (4,5)-bisphosphate targets double C2 domain protein B to the plasma membrane. Traffic 18:825–839. doi:10.1111/tra.12528

Mills SJ, Persson C, Cozier G, Thomas MP, Trésaugues L, Erneux C, Riley AM, Nordlund P, Potter BVL. 2012. A synthetic polyphosphoinositide headgroup surrogate in complex with SHIP2 provides a rationale for drug discovery. ACS Chem Biol 7:822–828. doi:10.1021/cb200494d

Milnik A, Heck A, Vogler C, Heinze HJ, de Quervain DJF, Papassotiropoulos A. 2012. Association of KIBRA with episodic and working memory: A meta-analysis. Am J Med Genet Part B Neuropsychiatr Genet 159 B:958–969. doi:10.1002/ajmg.b.32101

Nalefski EA, Falke JJ. 1996. The C2 domain calcium-binding motif: Structural and functional diversity. Protein Sci 5:2375–2390.

Nanda H, Heinrich F, Lösche M. 2015. Membrane association of the PTEN tumor suppressor: Neutron scattering and MD simulations reveal the structure of protein-membrane complexes. Methods 77:136–146. doi:10.1016/j.ymeth.2014.10.014

Naughton FB, Kalli AC, Sansom MSP. 2018. Modes of Interaction of Pleckstrin Homology Domains with Membranes: Toward a Computational Biochemistry of Membrane Recognition. J Mol Biol 430:372–388. doi:10.1016/j.jmb.2017.12.011

Naughton FB, Kalli AC, Sansom MSP. 2016. Association of Peripheral Membrane Proteins with Membranes: Free Energy of Binding of GRP1 PH Domain with Phosphatidylinositol Phosphate-Containing Model Bilayers. J Phys Chem Lett 7:1219–1224. doi:10.1021/acs.jpclett.6b00153

Pant S, Tajkhorshid E. 2020. Microscopic Characterization of GRP1 PH Domain Interaction With Anionic Membranes Shashank. 2Journal Comput Chem 41:489–499. doi:10.1002/jcc.26109.Microscopic

Parrinello M, Rahman A. 1981. Polymorphic transitions in single crystals: A new molecular dynamics method. J Appl Phys 52:7182–7190. doi:10.1063/1.328693

Posner MG, Upadhyay A, Ishima R, Kalli AC, Harris G, Kremerskothen J, Sansom MSP, Crennell SJ, Bagby S. 2018. Distinctive phosphoinositide- A nd Ca2+-binding properties of normal and cognitive performance-linked variant forms of KIBRA C2 domain. J Biol Chem 293:9335–9344. doi:10.1074/jbc.RA118.002279

Scott JL, Frick CT, Johnson KA, Liu H, Yong SS, Varney AG, Wiest O, Stahelin R V. 2020. Molecular analysis of membrane targeting by the C2 domain of the E3 ubiquitin ligase smurf1. Biomolecules 10:13–16. doi:10.3390/biom10020229

Shenoy S, Shekhar P, Heinrich F, Daou MC, Gericke A, Ross AH, Lösche M. 2012. Membrane association of the PTEN tumor suppressor: Molecular details of the protein-membrane complex from SPR binding studies and neutron reflection. PLoS One 7. doi:10.1371/journal.pone.0032591

Shenoy SS, Nanda H, Lösche M. 2012. Membrane association of the PTEN tumor suppressor: Electrostatic interaction with phosphatidylserine-containing bilayers and regulatory role of the C-terminal tail. J Struct Biol 180:394–408. doi:10.1016/j.jsb.2012.10.003

Slochower DR, Huwe PJ, Radhakrishnan R, Janmey PA. 2013. Quantum and all-atom molecular dynamics simulations of protonation and divalent ion binding to phosphatidylinositol 4,5-bisphosphate (PIP2). J Phys Chem B 117:8322–8329. doi:10.1021/jp401414y

Stahelin R V., Scott JL, Frick CT. 2014. Cellular and molecular interactions of phosphoinositides and peripheral proteins. Chem Phys Lipids 182:3–18. doi:10.1016/j.chemphyslip.2014.02.002

Südhof TC. 2004. The synaptic vesicle cycle. Annu Rev Neurosci 27:509–547. doi:10.1146/annurev.neuro.26.041002.131412

Treece BW, Heinrich F, Ramanathan A, Lösche M. 2020. Steering Molecular Dynamics Simulations of Membrane-Associated Proteins with Neutron Reflection Results. J Chem Theory Comput 16:3408–3419. doi:10.1021/acs.jctc.0c00136

Tribello GA, Bonomi M, Branduardi D, Camilloni C, Bussi G. 2014. PLUMED 2: New feathers for an old bird. Comput Phys Commun 185:604–613. doi:10.1016/j.cpc.2013.09.018

Tsuji T, Takatori S, Fujimoto T. 2019. Definition of phosphoinositide distribution in the nanoscale. Curr Opin Cell Biol 57:33–39. doi:10.1016/j.ceb.2018.10.008

Vermaas J V., Tajkhorshid E. 2017. Differential membrane binding mechanics of synaptotagmin isoforms observed in atomic detail. Biochemistry 56:281–293. doi:10.1021/acs.biochem.6b00468

Vickery ON, Stansfeld PJ. 2020. CG2AT. GitHub. doi:10.5281/zenodo.3890163

Wassenaar TA, Ingólfsson HI, Böckmann RA, Tieleman DP, Marrink SJ. 2015. Computational lipidomics with insane: A versatile tool for generating custom membranes for molecular simulations. J Chem Theory Comput 11:2144–2155. doi:10.1021/acs.jctc.5b00209

Wiesner S, Ogunjimi AA, Wang HR, Rotin D, Sicheri F, Wrana JL, Forman-Kay JD. 2007. Autoinhibition of the HECT-Type Ubiquitin Ligase Smurf2 through Its C2 Domain. Cell 130:651–662. doi:10.1016/j.cell.2007.06.050

Worby CA, Dixon JE. 2014. PTEN. Annu Rev Biochem 83:641–669. doi:10.1146/annurev-biochem-082411-113907

Yamamoto E, Kalli AC, Yasuoka K, Sansom MSP. 2016. Interactions of Pleckstrin Homology Domains with Membranes: Adding Back the Bilayer via High-Throughput Molecular Dynamics. Structure 24:1421–1431. doi:10.1016/j.str.2016.06.002

Zhang D, Aravind L. 2010. Identification of novel families and classification of the C2 domain superfamily elucidate the origin and evolution of membrane targeting activities in eukaryotes. Gene 469:18–30. doi:10.1016/j.gene.2010.08.006

Zhao H, Dupont J, Yakar S, Karas M, LeRoith D. 2004. PTEN inhibits cell proliferation and induces apoptosis by downregulating cell surface IGF-IR expression in prostate cancer cells. Oncogene 23:786–794. doi:10.1038/sj.onc.1207162

