## Supplementary material for "Binding of Ca^2+^-Independent C2 Domains to Lipid Membranes: a Multi-Scale Molecular Dynamics Study": SI

**SI Table S1. C2 Domain Distance vs. Time Exponential Decays.**

| Protein C2 | PC | | PC:PS | | $PC:PS:PIP_2$ | |
|---|---|---|---|---|---|---|
| | λ ($ps^{-1}$) | *B* (nm) | λ ($ps^{-1}$) | *B* (nm) | λ ($ps^{-1}$) | *B* (nm) |
| Smurf2 | 22 | 0.47 | 53 | 0.45 | 32 | 0.45 |
| RIM2 | 29 | 0.48 | 43 | 0.45 | 33 | 0.45 |
| KIBRA | 34 | 0.53 | 70 | 0.46 | 19 | 0.46 |
| PTEN | 15 | 0.46 | 13 | 0.56 | 19 | 0.45 |
| SHIP2 | 38 | 0.59 | 44 | 0.51 | 30 | 0.46 |
| PI3KCα | 23 | 0.48 | 68 | 0.45 | 44 | 0.45 |
| Mean ± SD | 27 ± 8 | 0.50 ± 0.05 | 48 ± 19 | 0.48 ± 0.04 | 30 ± 9 | 0.45 ± 0.01 |

The average decay curves derived from the protein-lipid minimum distance data in Fig. 3 were fitted with $D(t) = D_0 \exp(-\lambda t) + B$, where D is the average distance over the repeats, $D_0$ is the initial distance, $\lambda$ is the decay rate, $t$ is the simulation time, and $B$ is the average of the final minimum distance for all repeats. Mean and SD are mean and standard deviations for a specific lipid over all the different C2 domains. See SI Figure S1 for an example.

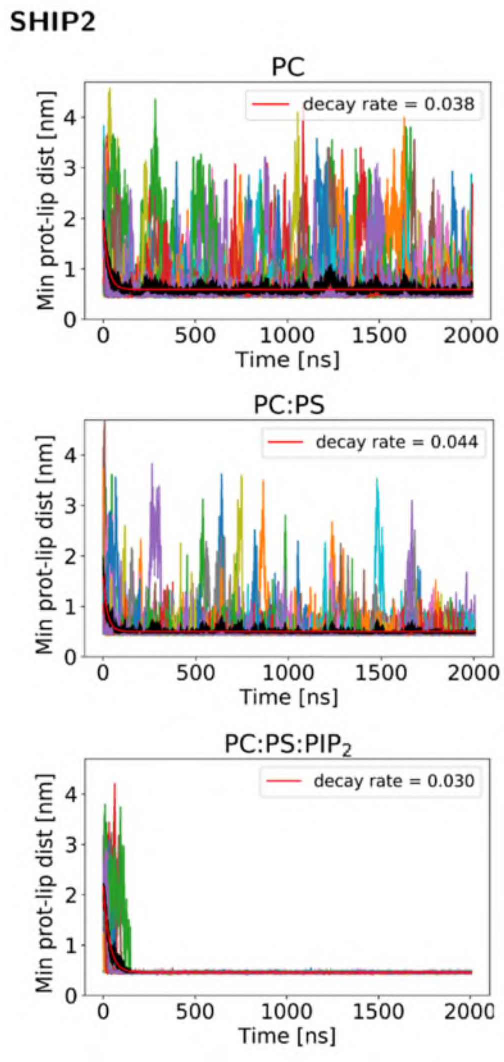

**SI Figure S1**

Minimum protein-lipid distances as a function of time for the SHIP2 C2 domain, fitted with exponential decay curves. The data are as shown in Fig. 3. See Table S1 for details of the decay curves (red) fitted to the averaged data (black line).

**A**

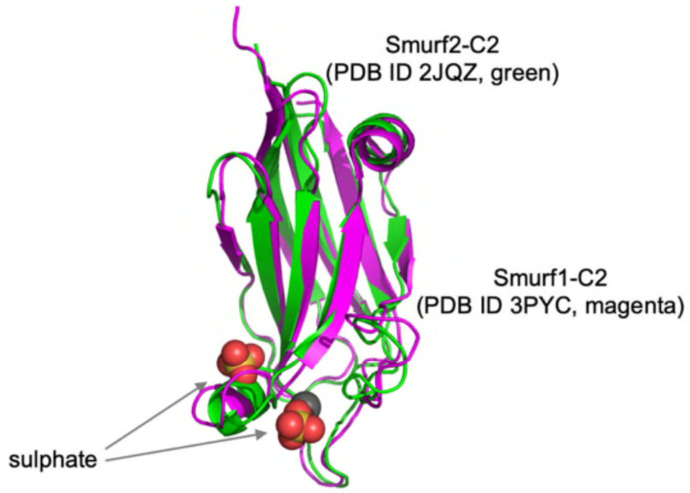

**B**

Alignment of 2JQZ (Smurf2-C2) and 3PYC (Smurf1-C2)

```
NW score     Identities            Positives      Gaps
615     116/133 (87%)        125/133        3/133 /2%)

2JQZ  1    GP--VKLRLTVLCAKNLVKKDFFRLPDPFAKVVVDGSGQCHSTDTVKNTLDPKWNQHYDL  58
           G   +K+RLTVLCAKNL KKDFFRLPDPFAK+VVDGSGQCHSTDTVKNTLDPKWNQHYDL
3PYC  1    GSEFIKIRLTVLCAKNLAKKDFFRLPDPFAKIVVDGSGQCHSTDTVKNTLDPKWNQHYDL  60

2JQZ  59   YIGKSDSVTISVWNHKKIHKKQGAGFLGCVRLLSNAINRLKDTGYQRLDLCKLGPNDNDT  118
           Y+GK+DS+TISVWNHKKIHKKQGAGFLGCVRLLSNAI+RLKDTGYQRLDLCKL P+D D
3PYC  61   YVGKTDSITISVWNHKKIHKKQGAGFLGCVRLLSNAISRLKDTGYQRLDLCKLNPSDTDA  120

2JQZ  119  VRGQIVVSLQSRD  131
           VRGQIVVSLQ+R
3PYC  121  VRGQIVVSLQTR   132
```

### SI Figure S2

**A** Smurf1-C2 (magenta, PDB ID 3PYC, including 2 sulphates) and Smurf2-C2 (green, PDB ID 2JQZ) aligned. **B** Sequence alignment of Smurf1-C2 and Smurf2-C2, made using BLAST (blast.ncbi.nlm.nih.gov).

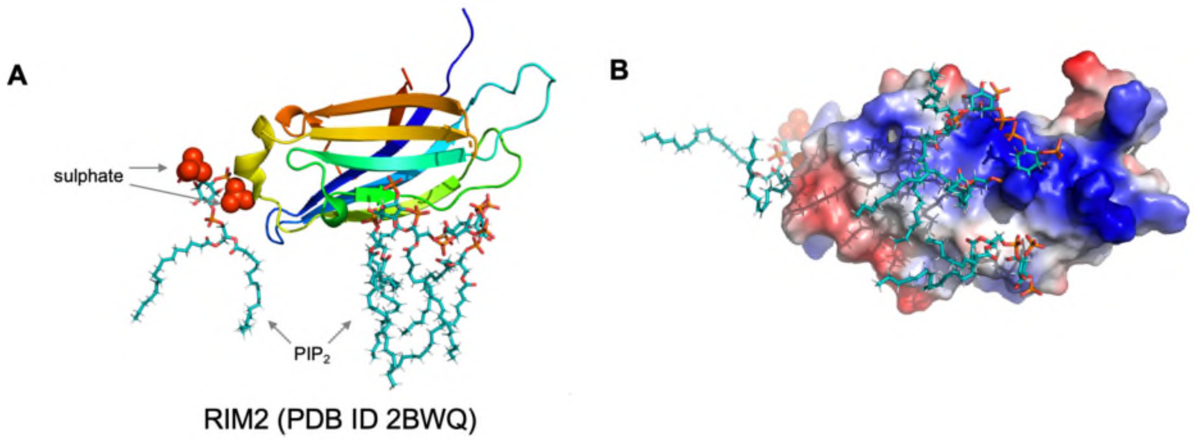

**SI Figure S3**

**A** Sulphates (in red/orange van der Waals format) in the crystal structure of RIM2 (PDB 2BWQ) are bound at the “bottom” of the C2 domain. The location of bound $PIP_2$ molecules (in ‘bonds’ format) from the atomistic simulation are also shown. **B** $PIP_2$ molecules observed bound in the simulation are shown alongside the surface of RIM2-C2 coloured according to its electrostatic potential (blue is positive charge and red is negative charge).

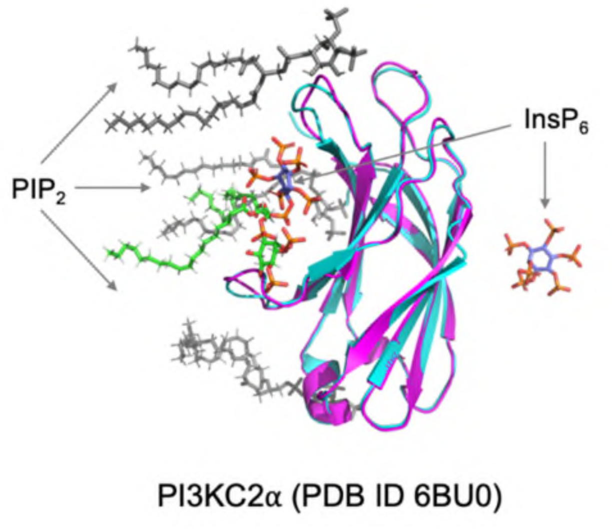

**SI Figure S4**

Crystal structure of PI3KC2α (PDB ID 6BU0; chain B; magenta) with 2 bound $InsP_6$ molecules, one at the front and one at the back. The simulated back-binding mode (cyan) structure is aligned with the crystal structure. One $PIP_2$ molecule (green) from the simulation was (in terms of headgroups) close to a bound $InsP_6$ molecule from the crystal (~0.85 nm) whereas the other $PIP2_s$ (gray) bound at other locations.
